# Endoplasmic reticulum stress signaling actively contributes to therapy resistance in colorectal cancer

**DOI:** 10.1101/2024.01.07.574523

**Authors:** Daisuke Sasaki, Natsuki Sato, Dirk Wilhelm, Julius Fischer, Julia Gissibl, Masatoshi Nakatsuji, Dirk Haller, Hideki Ishihara, Klaus-Peter Janssen

## Abstract

**Purpose:** We investigated the involvement of endoplasmic reticulum (ER) stress signaling in cancer cell responses to chemo- and radiotherapy, focusing on three main ER stress mediators, the transcription factors ATF4, XBP1 and ATF6.

**Methods:** Public cancer genome datasets were assessed for alterations in ER stress mediators. Surgically resected colorectal cancer tissues were tested by flow cytometry and used to generate patient-derived organoids. Human cell lines and organoids were characterized under oxaliplatin treatment, alone or combined with pharmacological inhibitors of the three ER stress branches, or X-ray irradiation, for cytotoxicity, activation of ER stress and proteome changes. To monitor ER stress in real time, stable HEK293 kidney epithelial cell lines were established expressing ATF4, XBP1, or ATF6, fused with a fluorophore.

**Results:** Genomic amplification of ATF6, but not ATF4 or XBP1, was frequent in solid tumor entities like breast, lung and colorectal cancer and significantly associated with reduced disease-free survival. In colorectal cancer, increased ATF6 was associated with genetic instability. Basal ER stress mediator expression was correlated to chemoresistance in colorectal cancer cell lines, and generally high in cancer cells compared to HEK293 cells. With proteomics and live HEK293-based reporter lines, we noted that oxaliplatin treatment induced ER stress in a remarkably different way from the canonical ER stress inducer thapsigargin. Moreover, modulation of ER stress signaling by exogenous expression of the stress mediators positively affects chemoresistance, and pharmacological inhibition of ATF6 sensitizes ER-stressed HCT116 colorectal cancer cells to chemotherapy. Of note, cellular stress responses was strongly dependent on the individual transcription factor: XBP1-driven response appeared multi-functional, involved in ribosome biogenesis stress and associated with oxaliplatin resistance. ATF6-dependent stress signaling was involved in DNA damage repair, and was essential for radioresistance. Moreover, chemoresistance in HCT116 cancer cells was impaired by pharmacological ATF6 inhibition.

**Conclusion:** Activation of the ER stress signaling may be critically involved in acquired chemo- and radioresistance. Due to their apparent cytoprotective roles, ATF6 and XBP1 could be attractive predictive biomarkers and putative therapeutic targets.

**SUMMARY:** To address their roles in the clinical context, genomic alterations of ATF4, XBP1 and/or ATF6 in human solid tumors were assessed with respect to prognosis and genomic instability. Moreover, surgically resected CRC patient tissues were tested for expression of ER stress markers by flow cytometry and associated with clinical characteristics. In addition, a panel of human cell lines and patient-derived colon organoids were characterized under therapeutic conditions for expression and activation of ER stress proteins, and resulting cytotoxicity was determined. To monitor and modulate ER stress activation in live cells with subcellular resolution, stable reporter cell lines expressing ATF4, sXBP1 or ATF6 proteins fused with a fluorophore were established. These lines were tested for gene or protein expression and cytotoxicity assays to analyze how activation or inhibition of ER stress proteins affects the cellular responses to oxaliplatin treatment or X-ray irradiation. Finally, mass spectrometric proteome analysis was performed to obtain an unbiased readout on the cellular responses to chemotherapy driven by the activation of the ER stress proteins.

## Introduction

In addition to surgery, chemotherapy and radiotherapy are still by large the main treatment options for cancer patients worldwide. However, limited primary response rates and disease recurrence remain challenging. Here, we focused on colorectal cancer (CRC) as third most prevalent malignancy and second most lethal cancer worldwide (Sung et al., 2021). Although the primary treatment option for CRC is surgical resection, nearly one out of four patients already presents with unresectable metastases when diagnosed, and around 30% of patients who have undergone curative-intent resection will eventually develop disease recurrence (van der Stok et al., 2017). For such patients, chemotherapy (radio-chemotherapy for rectal cancer) is the leading strategy to control disease by pre-operative reduction of the tumor and metastasis size, and suppression of further progression and spread. Oxaliplatin (OxaPt), a third-generation platinum agent, is a crucial component of standard chemotherapy for advanced CRC (de Gramont et al., 2000). OxaPt-based chemotherapy has demonstrated improved efficacy compared to other regimens, significantly reducing recurrence and mortality (André et al., 2015; Souglakos et al., 2006). However, intrinsic or acquired resistance to OxaPt is a central challenge in the CRC treatment, and the underlying mechanisms remain to be elucidated (Douillard et al., 2013; Martinez-Balibrea et al., 2015).

One proposed mechanism for therapy resistance is through cellular stress-response programs, among them adaptive pathways that ensure the homeostatic function of the endoplasmic reticulum (ER), collectively called unfolded protein response (UPR), or ER stress response (Kaufman, 2002; Mori, 2000). ER stress can be triggered by a variety of cell-intrinsic (e.g., aneuploidy) or environmental factors that cause an overload of unfolded proteins in the ER thereby activating signals to either restore ER homeostasis or eliminate irreversibly damaged cells by apoptosis (Hetz et al., 2015). The ER stress response is initiated by three major ER-resident receptors — protein kinase RNA-like ER kinase (PERK), inositol-requiring gene 1 (IRE1) and activating transcription factor 6 (ATF6) (Harding et al., 1999; Haze et al., 1999; Shamu & Walter, 1996). Activated PERK phosphorylates the eukaryotic translation initiation factor 2α (eIF2α) to induce global translational repression and selective translation of transcription factor ATF4, which regulates the expression of a gene cluster responsible for amino acid transport, redox control, autophagy and apoptotic pathways as well as protein folding (Harding et al., 2003; Palam et al., 2011; Vattem & Wek, 2004). Active IRE1 catalyzes mRNA splicing of the X-box binding protein 1 (XBP1), resulting in the shift of the reading frame that translates to the transcription factor sXBP1 (spliced isoform of XBP1). In turn, sXBP1 modulates the expression of genes involved in a wide range of proteostatic processes including ER folding of proteins, the ER-associated degradation (ERAD) of unfolded proteins, lipid metabolism, secretory pathways, glucose homeostasis and inflammatory responses (Acosta-Alvear et al., 2007; Byrd & Brewer, 2012; Calfon et al., 2002; Lee et al., 2003; Wu et al., 2015). IRE1 can also degrade other RNAs through a process named regulated IRE1-dependent decay (RIDD) for further translational regulation (Hollien et al., 2009). ATF6 translocates to the Golgi apparatus under ER stress conditions and is processed into an N-terminal fragment (nATF6), which serves as a transcription factor, regulating genes involved in ER folding, modification of proteins and ERAD (Kokame et al., 2001; Roy & Lee, 1999; Yamamoto et al., 2007). Of note, besides these canonical proteostatic signaling pathways, functional links between the ER stress response and the DNA damage response (DDR) were suggested, where those transcription factors promote the expression of genes for the DNA damage recognition and repairing (González-Quiroz et al., 2020). Overall, the ER stress response constitutes a central hub that integrates various adaptive pathways for survival from proteotoxic and genotoxic stresses.

We and others have previously documented the involvement of UPR and ER stress in human pathologies such as IBD and cancer (Haller papers). Moreover, micriobial triggers are well established as inducers of ER stress for the ATF6 branch (Coleman et al., 2018).

In cancer cells, the ER stress signaling is basally activated to cope with not only external hostile conditions such as hypoxia, acidosis and nutrient deprivation but also intrinsic proteotoxic stresses such as genomic instability, oncogene activation and increased glycolysis (Dejeans et al., 2014; Moenner et al., 2007). Recent studies have shown that the activation of the ER stress signaling promotes resistance to chemotherapy and might also contribute to the adaptive survival pathways during radiotherapy in cancer cells (Bahar et al., 2019; Kim et al., 2019). However, it is poorly understood how these three branches are involved in therapy resistance of cancer cells, and therefore, further experimental evidence will be needed to explore their potential as predictive biomarkers and therapeutic targets.

In this study, we aimed to better understand the role of the ER stress pathways in cellular responses to chemo- and radiotherapy. We focused on the nuclear activity of the transcription factors ATF4, sXBP1 and nATF6 and provide insights into their individual role in confering resistance to cancer therapy, which might offer new avenues for patient stratification and therapeutic intervention.

## Materials and methods

### Clinical samples and data

Human colorectal tissues were obtained from CRC patients undergoing surgical interventions at the Dept. of Surgery, Klinikum Rechts der Isar, Technical University of Munich (Munich, Germany), with written informed consent and prior approval by the ethics committee at the Faculty of Medicine of TUM, following guidelines and regulations proposed in the Declaration of Helsinki (330/18S; 2022-169-S-KH). A human cancer genomic dataset from The Cancer Genome Atlas (TGCA), consisting of 10,953 patients in the Pan-Cancer Atlas Studies, was analyzed using the cBioPortal platform (https://www.cbioportal.org) for low/high levels of gene-level amplification of ATF4, XBP1 and ATF6. A pan-cancer dataset (n=10,953 patients), and a CRC subset (n=594 patients) were analyzed for the ATF6 genomic amplification combined with point mutations and/or gene-level homozygous deletions in the TP53 gene (MUT/HOMDEL) in association with the aneuploidy score and prognosis.

### Cell culture

Human cell lines were obtained from the American Type Culture Collection (Manassas, VA, USA) and maintained at 37 °C in a humidified 7% CO_2_ atmosphere. DLD1, HCT116, HEK293, HT29, SW480 and SW620 were cultured in DMEM (Thermo Fisher Scientific, Waltham, MA, USA) with 7% fetal calf serum (FCS), 2 mM L-glutamine, 50 U/ml penicillin and 50 μg/ml, streptomycin. HepaRG cells were cultured in WEM (Thermo Fisher Scientific) with 10% FCS, 2 mM GlutaMAX (Thermo Fisher Scientific), 5 ng/ml human insulin (Sanofi, Paris, France), 50 μM hydrocortisone hemisuccinate (Sigma-Aldrich, St. Louis, MO, USA), 10 U/ml penicillin and 10 μg/ml streptomycin.

### Generation of stable transfected cell clones

HEK293 cells were cultured in 6-well plates and then transfected with 4 μg of previously described plasmid vectors encoding ER stress reporter genes ATF4-mS (#115970, Addgene, Cambridge, MA, USA), XBP1-mN (#115971), ATF6-GFP (#32955) or H2A-mS (#85051) by calcium phosphate precipitation. The media were refreshed 24 h after transfection. Two days post transfection, the cells were subjected to antibiotic selection with 200 μg/ml Hygromycin B (Invitrogen, Carlsbad, CA, USA) or 400 μg/ml G418 (Thermo Fisher Scientific). Resistant colonies were selected and subcloned by limiting dilution. Positive cells were further enriched by FACS (Aria III, BD Biosciences, San Jose, USA).

### Primary human colorectal organoids

Crypts were isolated from minced mucosa of surgically reseceted human non-diseased colon tissue by treating with 5 mM EDTA in PBS and mechanical dissociation. The crypts were embedded in Matrigel (Corning, Corning, NY, USA) and cultured with the Advanced DMEM/F-12 (Thermo Fisher Scientific) containing 10 mM HEPES, 2 mM GlutaMAX, 10 mM nicotinamide, 1 mM N-acetylcysteine, B-27 supplement (Thermo Fisher Scientific), 10 μM Y-27632 (Sigma-Aldrich), 50 ng/ml EGF (PeproTech, Rocky Hill, NJ, USA), 50 nM A83-01 (Tocris Bioscience, Bristol, UK), 10 nM PGE2 (Sigma-Aldrich), 7.5 μM SB202190 (Sigma-Aldrich), 50% L-WRN conditioned medium, Antibiotic-Antimycotic (Thermo Fisher Scientific) and 100 μg/ml normocin at 37 °C in a humidified 7% CO_2_ atmosphere.

### Chemical and radiation treatment

Thapsigargin, Tg (Sigma-Aldrich), was used for canonical ER stress induction according to published data, dissolved in DMSO at 10 mM concentration and stored at –20 °C. (Qui et al., 2011). Oxaliplatin (OxaPt) was supplied as a 5 mg/ml solution by the Pharmacy of Klinikum Rechts der Isar, TUM, and stored at 4 °C. Compounds GSK2656157, STF-083010 and Ceapin-A7 (Sigma-Aldrich) were dissolved in DMSO at 5 mg/ml concentration and stored at –20 °C. The reagents were diluted in the culture medium to achieve the final concentrations. DMSO (H_2_O for OxaPt) was used as the corresponding vehicle control for the treatment. Irradiation was performed using an X-ray device (RS225, Gulmay Medical, Surrey, UK; 200 keV, 15 mA).

### Flow cytometry

The clinical tissue specimens were digested into single cells with 0.5 mg/ml Collagenase Type IV (Thermo Fisher Scientific) in DMEM containing 25 mM HEPES, followed by filtration at 40 µm. Cells were fixed with 3% PFA, quenched by 50 mM ammonium chloride after 20 min. The cells were permeabilized with 0.1% Triton X-100, blocked with 2% BSA and incubated with primary rabbit anti-XBP1 antibody (ab37152, 1:200, Abcam, Cambridge, MA, USA), ATF6 antibody (ab203119, 1:200, Abcam) or rabbit IgG isotype-matched control (ab37415, Abcam) in 0.5% BSA at room temperature for 20 min. Subsequently, cells were stained with secondary goat anti-rabbit Alexa488 (1:500, Thermo Fisher Scientific) and DAPI (2 μg/ml) in 0.5% BSA at room temperature for 10 min. 2D-cultured cells were trypsinized before analysis. Apoptosis detection was performed by staining with Annexin V-FITC and propidium (PI), using a commercial kit (Abcam). Fluorescence was detected by a CytoFLEX device (Beckmann Coulter, Brea, CA, USA) and analyzed using FlowJo software (v.10.8.1, BD Biosciences). Signal positivity was determined with the cut offs at 95 percentiles of isotype-matched negative control, or untreated reporter cells.

### Cell viability assay

The cell viability was determined by the XTT or MTT assay. For the XTT assay, cells were seeded in 96-well plates (1 × 10^3^ cells/well) one day before the start of treatment (in triplicate for each condition). After 24 h of treatment, cell viability was evaluated using the XTT Cell Proliferation Assay Kit II (Roche Diagnostics, Basel, Switzerland) according to the manufacturer’s protocol. For the MTT assay, cells were seeded in 96-well plates (2 × 10^3^ cells/well) and analyzed with the MTT Cell Growth Assay Kit II (Sigma-Aldrich) according to the manufacturer’s instruction. The cell proliferation index was calculated as change of absorbance to untreated conditions.

### Colony formation assay

Cells were seeded at a density of 400 or 800 cells/well in 12-well plates one day before treatment (in triplicate for each condition). After irradiation, the cells were cultured for 1–2 weeks until visible colonies formed. The colonies were fixed with 3% PFA and stained with 0.1% crystal violet (Sigma-Aldrich). Colonization was then measured as a ratio of areas occupied by colonies to the total surface areas using the Fiji/ImageJ software (v.2.3.0, NIH, Bethesda, MD, USA). Cell survival rate was calculated by comparing colonization to untreated conditions.

### Immunofluorescence

Cells were seeded on coverslips precoated with 0.1% gelatin two days before the start of treatment. At the time points after 0, 0.5, 1, 3, 6 and 24 h of treatment, the cells were fixed with 3% PFA and quenched by treatment with 50 mM ammonium chloride after 20 min. Then, the cells were permeabilized with 0.1% Triton X-100, followed by blocking with 2% BSA. The cells were incubated with primary anti-Golgin-97 antibody (A-21270, 1:100, Thermo Fischer Scientific) in 2% BSA at 4 °C overnight, subsequently stained with secondary goat anti-mouse Cy3 or anti-mouse Alexa488 (1:300, Thermo Fischer Scientific) and DAPI (2 μg/ml) in 2% BSA at room temperature for 2 h. The coverslips were mounted on glass slides with glycerol gelatin (Sigma-Aldrich). Optical section images were obtained with a 100 × oil immersion objective using the AxioObserver Z1 microscope equipped with an ApoTome system (Carl Zeiss, Stuttgart, Germany).

Deconvolution and image processing were performed using the ZEN software (v.3.0, Carl Zeiss). The colocalization of different fluorescent probes was analyzed by the Fiji/ImageJ software. For dead cell quantification in organoids, the culture was washed with TBS and stained with SYTOX-Green (1:30,000, Thermo Fisher Scientific) and Hoechst-33342 (1:1000, Thermo Fisher Scientific) for 20 min and imaged using the AxioObserver Z1 microscope (Carl Zeiss). Fluorescence from individual organoids was quantified by the Fiji/ImageJ software.

### qRT-PCR

The RNeasy Mini Kit (Qiagen, Hilden, Germany) was used for the extraction of total RNA from cells. The concentration and purity of the RNA samples were measured by NanoDrop 1000 (Thermo Fisher Scientific). RNA was reverse transcribed into cDNA using the RevertAid RT Kit (Thermo Fisher Scientific). Gene expression was detected using the LightCycler 480 SYBR Green I Master Mix or Probes Master Mix kit (Roche Diagnostics) after PCR (5 min at 95 °C; 45 cycles of 10 s at 95 °C, 10 s at 60 °C, and 10 s at 72 °C). The expression of mRNA was compared to HPRT and quantified using the 2^− ΔΔCt^ method. The signals were normalized based on untreated HEK293/HCT116 cells. The primers were: ATF4 forward, 5’-TCT CCA GCG ACA AGG CTA A-3’; ATF4 reverse, 5’-CCA ATC TGT CCC GGA GAA-3’; XBP1 forward, 5’-CCC TGG TTG CTG AAG AGG-3’; XBP1 reverse, 5’-TGG AGG GGT GAC AAC TGG-3’; sXBP1 forward, 5’-GCT GAG TCC GCA GCA GGT-3’; sXBP1 reverse, 5’-CAG ACT ACG TGC ACC TCT GC-3’, uXBP1 forward, 5’-CTG GGT CCA AGT TGT CCA GAA T-3’, uXBP1 reverse, 5’-CAG ACT ACG TGC ACC TCT GC-3’, ATF6 forward, 5’-AAT ACT GAA CTA TGG ACC TAT GAG CA-3’; ATF6 reverse, 5’-TTG CAG GGC TCA CAC TAG G-3’; HPRT forward 5’-GAC CAG TCA ACA GGG GAC AT-3’, HPRT reverse 5’-GTG TCA ATT ATA TCT TCC ACA ATC AAG-3’. The primers for s/uXBP1 were adopted from a previous study (Yoon et al., 2019).

### XBP1 mRNA splicing assay

cDNA (1 μg/sample) was amplified by PCR (3 min at 94 °C; 40 cycles of 30 s at 94 °C, 30 s at 60 °C, and 60 s at 72 °C) using the Taq DNA Polymerase kit (Thermo Fischer Scientific) with primers for human XBP1. The products were resolved on a 2.5% agarose gel with EtBr detection, and signals were quantified using the Fiji/ImageJ software.

### Western blotting

Cells were lysed with RIPA buffer containing 0.1% SDS and protease/phosphatase inhibitors: 1 mM Na_3_VO_4_, 1 mM NaF, 1 mM Pefabloc, Protease Inhibitor Cocktail (Roche Diagnostics), 1 mM β-glycerophosphate, 1 mM benzamidine, 1 mM PMSF, 1 μg/ml Pepstatin A. Proteins were boiled in Laemmli buffer with 100 mM DTT at 95 °C for 5 min before SDS-PAGE. The gel was initially run at 80 V and switched to 100 V after bromophenol blue entered the resolving gel. The proteins were blotted to nitrocellulose membranes (12 V, 1 h). The membranes were rinsed with PBST and blocked with 5% skim milk (Carl Roth, Karlsruhe, Germany) in PBST. The membranes were incubated with the following primary antibodies in 5% skim milk at 4 °C overnight: rabbit anti-ATF4 antibody (ab270980, 1:1000, Abcam), XBP1 antibody (ab37152, 1:500, Abcam), ATF6 antibody (ab203119, 1:500, Abcam), p-eIF2α antibody (#9721, 1:1000, Cell Signaling Technology, Boston, MA, USA), BiP antibody (#3177, 1:1000, Cell Signaling Technology), mouse anti-β-actin antibody (#3700, 1:2000, Cell Signaling Technology), PARP1 antibody (sc74470, 1:1000, Santa Cruz Biotechnology, Dallas, TX, USA). The membranes were rinsed with PBST and subsequently incubated with secondary goat anti-rabbit-HRP or anti-mouse-HRP antibody (1:4000, Jackson ImmunoResearch, West Grove, USA) in 5% skim milk at room temperature for 1 h. The membranes were rinsed with PBST and subjected to chemiluminescent protein detection with the ECL Western Blotting Substrate (Thermo Fischer Scientific).

### Cell fractionation

Cells were thoroughly resuspended by pipetting in ice-cold 0.3% NP-40/PBS to lyse cytoplasmic membranes. One-third volume of the suspension was transferred to a reaction tube as a “whole cell lysate (W)” fraction. The remaining suspension was centrifuged at 15000 rpm, 4 °C for 1 min, and half the volume of the supernatant was transferred to a new reaction tube as a “cytoplasmic (C)” fraction. The remaining supernatant was discarded, and the pellet was washed three times with 0.3% NP-40/PBS by pipetting and centrifugation at 15000 rpm, 4 °C for 1 min. The pellet was resuspended in 0.3% NP-40/PBS (half the volume of the other fractions) to obtain a “nuclear (N)” fraction. These fractions were boiled with Laemmli buffer and 100 mM DTT at 95 °C for 5 min and used for SDS-PAGE and immunoblotting.

### Proteomics

Cells were lysed by incubating at 95°C for 10 min with agitation in a previously reported lysis buffer (Kulak et al., 2014). The extracted proteins were digested with trypsin and Lys-C endopeptidase at a ratio of 1:100 to the protein amount (37°C, overnight). The following day, the trypsinization was quenched by adding 200 µl 1% trifluoroacetic acid (TFA) in isopropanol. Peptides were then desalted using styrene divinylbenzene-reverse phase sulfonate Stage-tips (Merck, Darmstadt, Germany) according to the manufacturer’s protocol. The peptides were resuspended in 20 µl buffer A (2% acetonitrile, 0.2% TFA) and analyzed in single shots in the timsTOF HT mass spectrometer (Bruker Daltonics, Billerica, MA, USA) coupled to a Vanquish Neo liquid chromatography system (Thermo Fisher Scientific): 500 ng of peptides were loaded on a 45 cm HPLC column packed with 1.9 µM C18 ReproSil Pure-AQ particles (Dr. Maisch, Ammersbach, Germany) and eluted over a 100 min gradient from 7% to 38% buffer B in 90 min followed by an increase to 98% in 3 min, and a column wash at 98% buffer B for 7 min (buffer A: 0.1% formic acid in LC-MS grade H_2_O; buffer B: 0.1% formic acid, 80% acetonitrile, and 19.9% LC-MS grade H_2_O) with a flow rate of 350 nl/min.

MS data were acquired in a data-independent mode with parallel accumulation followed by serial fragmentation (dia-PASEF) as previously described (Meier et al., 2015). Mass spectra were acquired from 100 to 1700 m/z and 0.7 to 1.45 1/K0 with a 100 ms ramp time at 100% duty cycle at a 1.06 s cycle time. The DIA-NN software (v.1.8.1, https://github.com/vdemichev/DiaNN/) was used to calculate label-free intensities in the library-free mode. Oxidized methionine and carbamidomethyl was set as variable and fixed modification, respectively. FDR (also referred to as *q*-value) was set at 0.01 for both peptide and protein levels. The minimal length of the peptides considered for identification was set at seven amino acids, and the option “match between runs” was enabled. Quantification mode was set to “Robust LC (high precision)” and all other settings were left default. The UniProt database from humans (*Homo sapiens*) was used for peptide and protein identification.

### Statistical analysis

Pairwise comparison between two groups was performed by Student t-test or Mann-Whitney tests using the Prism software (v.9.5.1, GraphPad, La Jolla, CA, USA), and data are presented as mean (± standard deviation). Comparison of survival rates was performed by Kaplan-Meier analysis with log-rank tests. Differences with *p, q* ≤ 0.05 were considered to be significant. In bioinformatics of the proteome data, a customized Python-based tool was used for statistical analysis and data visualization. Complementarily, the R environment (v.4.2.2, R Foundation for Statistical Computing, Vienna, Austria) was used for data analysis and representation. The Fisher’s exact test was used to generate significant enrichment gene ontology (GO) terms with an FDR cut-off of 0.05.

## Results

### ATF6 is frequently amplified in human cancer and associated with decreased survival, as well as with genomic instability in colorectal cancer

To address the involvement of ER stress signaling in cancer formation and therapy response, a publicly available dataset from the Pan-Cancer Atlas in the cBioPortal database was analyzed. We focused on the frequency of genomic amplification of ATF4, XBP1 and ATF6 and their association with disease-free survival (Fig. 1A). ATF6 was the most frequently amplified in major human solid cancer entities (Fig. 1A, top left). Notably, the ATF6 gene amplification was almost exclusive, compared to the other genes, in CRC. Kaplan-Meier analysis revealed that ATF6 gene amplification was significantly associated with reduced disease-free survival (*p* ≤ 0.0001), while the amplification of ATF4 and XBP was not (*p* = 0.2232 and 0.00542, respectively). All three genes showed a significant decrease of overall survival when upregulated at the genome level (*p* ≤ 0.0006, Fig. S1A).

**Fig. 1.**
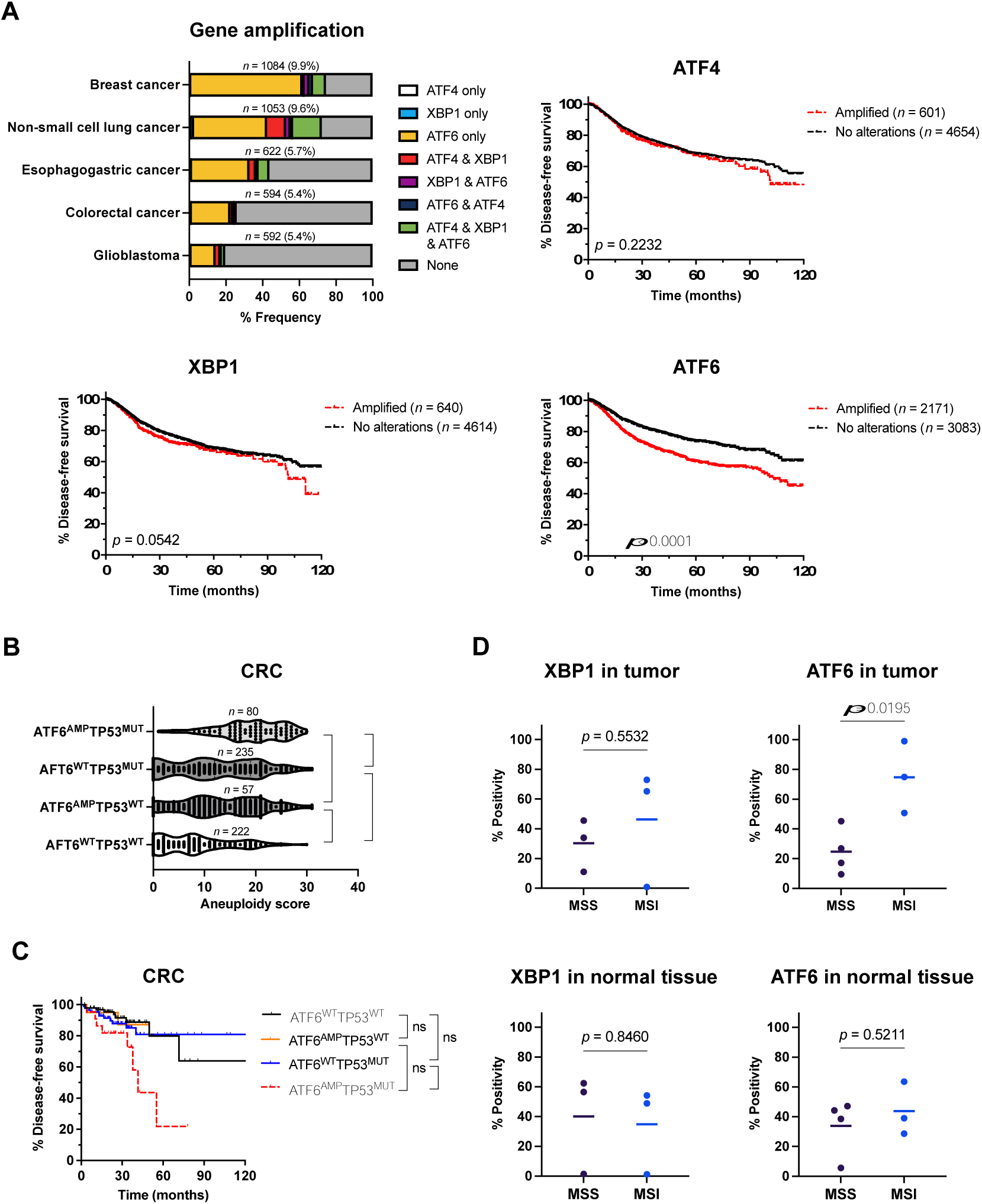
Amplification of ATF6 specifically associates with genomic instability and disease-free survival in public genomic datasets, and in surgically resected human CRC. **(A)** Frequencies of ATF4, XBP1 and/or ATF6 gene amplification in patients with major solid cancer types (left top) and Kaplan-Meier analysis of disease-free survival among cancer patients with amplified ATF4, XBP1 or ATF6 compared to patients with no alterations from the public TCGA datasets. **(B)** Comparison of aneuploidy scores between CRC patients with ATF6 gene amplification (ATF6^AMP^) plus/minus TP53 mutation or homozygous deletion (TP53^MUT^) and patients with no alterations (TP53^WT^ and/or ATF6^WT^). \**p* ≤ 0.05, \*\**p* ≤ 0.01, \*\*\*\**p* ≤ 0.0001. **(C)** Kaplan-Meier analysis of disease-free survival among CRC patients present with TP53^MUT^ and/or ATF6^AMP^ signatures, compared to patients with no alterations (TP53^WT^ and/or ATF6^WT^) from the public TCGA datasets. \**p* ≤ 0.05, \*\**p* ≤ 0.01, \*\*\*\**p* ≤ 0.0001. **(D)** The positivity of XBP1 and ATF6 was assessed by flow cytometry and compared between MSI group (*n* = 3) and MSS group (*n* = 3–4) in colon tumor and adjacent normal colon tissues. MSI, microsatellite instability. MSI, microsatellite instable; MSS, microsatellite stable or below the detection limit.

Moreover, we found a close relationship of genomic ATF6 amplification with genetic instability in colorectal cancer (Fig. 1B, C; Fig. S1B, S1C). Cancer patients were categorized into subsets without alterations in the TP53 gene (TP53^WT^), or with TP53 mutations and/or homozygous deletions (TP53^MUT^), a genomic constellation known to be associated with aneuploidy in cancer in a TCGA dataset (Taylor et al., 2018). First, these patient subgroups were compared based on the aneuploidy score, as defined in the cBioPortal platform by the number of chromosome arms with arm-level copy-number alterations (Fig. 1B; Fig. S1B). Patients with genomically amplified ATF6 exhibited a significant increase in aneuploidy score (*p* ≤ 0.0001) compared to those with normal ATF6 and TP53 status (ATF6^WT^TP53^WT^), even if TP53 was unaltered (ATF6^AMP^-TP53^WT^). As expected, TP53 genomic alterations alone were significantly associated with increased aneuploidy (*p* ≤ 0.0001, ATF6^WT^-TP53^MUT^ vs. ATF6^WT^-TP53^WT^). Of note, patients with ATF6 genomic amplification, combined with TP53 alterations (ATF6^AMP^-TP53^MUT^) showed the most substantial increase in aneuploidy scores (*p* ≤ 0.0001) compared to single alterations either in ATF6 or TP53 (ATF6^AMP^-TP53^WT^ and ATF6^WT^-TP53^MUT^).

Finally, potential prognostic differences between these subgroups were assessed using Kaplan-Meier analysis (Fig. 1C; S1C). In the CRC dataset, possibly due to relatively small cohort sizes, no significant difference was observed in overall survival among these groups. However, the disease-free survival of the ATF6^AMP^TP53^MUT^ co-occurrence group significantly decreased compared to that of the ATF6^WT^TP53^MUT^ group. Neither ATF6 amplification nor TP53 alterations individually showed significant association with changes in disease-free survival (Supplementary Fig. xx). In the Pan-Cancer dataset, ATF6^WT^TP53^WT^ showed the most favorable survival compared to the other groups. Genomic ATF6 amplification, even in the absence of TP53 mutations (ATF6^AMP^TP53^WT^), was significantly associated with reduced overall and disease-free survival (*p* ≤ 0.01). In addition, combined alteration of both ATF6 and TP53 (ATF6^AMP^TP53^MUT^) was associated with even further decreased survival (Fig. S1C). Notably, alteration of TP53 was not associated with worse overall survival in cases with amplified ATF6, since the group with compound mutations (ATF6^AMP^TP53^MUT^) did not differ significantly from patients with only amplified ATF6 (ATF6^AMP^TP53^WT^) (Fig. S1C, left).

Furthermore, we analyzed the expression of XBP1 and ATF6 on protein levels in a small cohort of surgically resected human CRC using flow cytometry (*n* = 7), compared to clinical characteristics (summarized in Table S1). Three cases were diagnosed with high microsatellite instability (MSI-high), indicating mismatch-repair deficiency, a known predictor of chemotherapy response (Cherri et al., 2022). Interestingly, ATF6 positivity in tumors was significantly associated with MSI-high status, while ATF6 levels in adjacent normal colon and XBP1 in tumors remained essentially unchanged (Fig. 1D, *p* = 0.0195). ATF6 levels in MSI-high tumors tended to be higher than in matched normal colon, although the difference was not significant (Fig. S1D, *p* = 0.1695). Regarding association with clinical parameters, tumors in earlier T-stages had significantly higher ATF6 positivity than more advanced T-stages (Table S1 and Fig. S1E, *p* = 0.0381). No further significant correlations were observed between ATF6 or XBP1 staining and clinical or pathological features such as staging, grading, anatomical tumor location, pN-stage, or pM-stage (Table S1, *p* > 0.05).

### Basal ER stress activity and chemoresistance in human cell lines

Next, we examined baseline ER stress activity in established human cell lines and their association with sensitivity against the chemotherapeutic agent oxaliplatin (OxaPt). A range of human CRC lines and control cell lines from kidney and liver were analyzed by qRT-PCR to assess basal gene expression of ATF4, sXBP1 and uXBP1 (spliced and unspliced isoforms of XBP1, respectively) and ATF6 (Fig. 2A). Western blotting of subcellular fractions from HCT116, HT29 and DLD1 cells confirmed nuclear localization of sXBP1 and nATF6 (nuclear proteolytic fragment of ATF6), indicating constitutive ER stress signaling (Fig. S2A). We observed a positive, but not significant, correlation between basal ER stress gene expression levels and OxaPt resistance, based on IC_50_ values obtained by cell proliferation assays (Fig. S2B–C, *n* = 6, *r* = 0.3903–0.7362, *p* = 0.0952–0.4442).

**Fig. 2.**
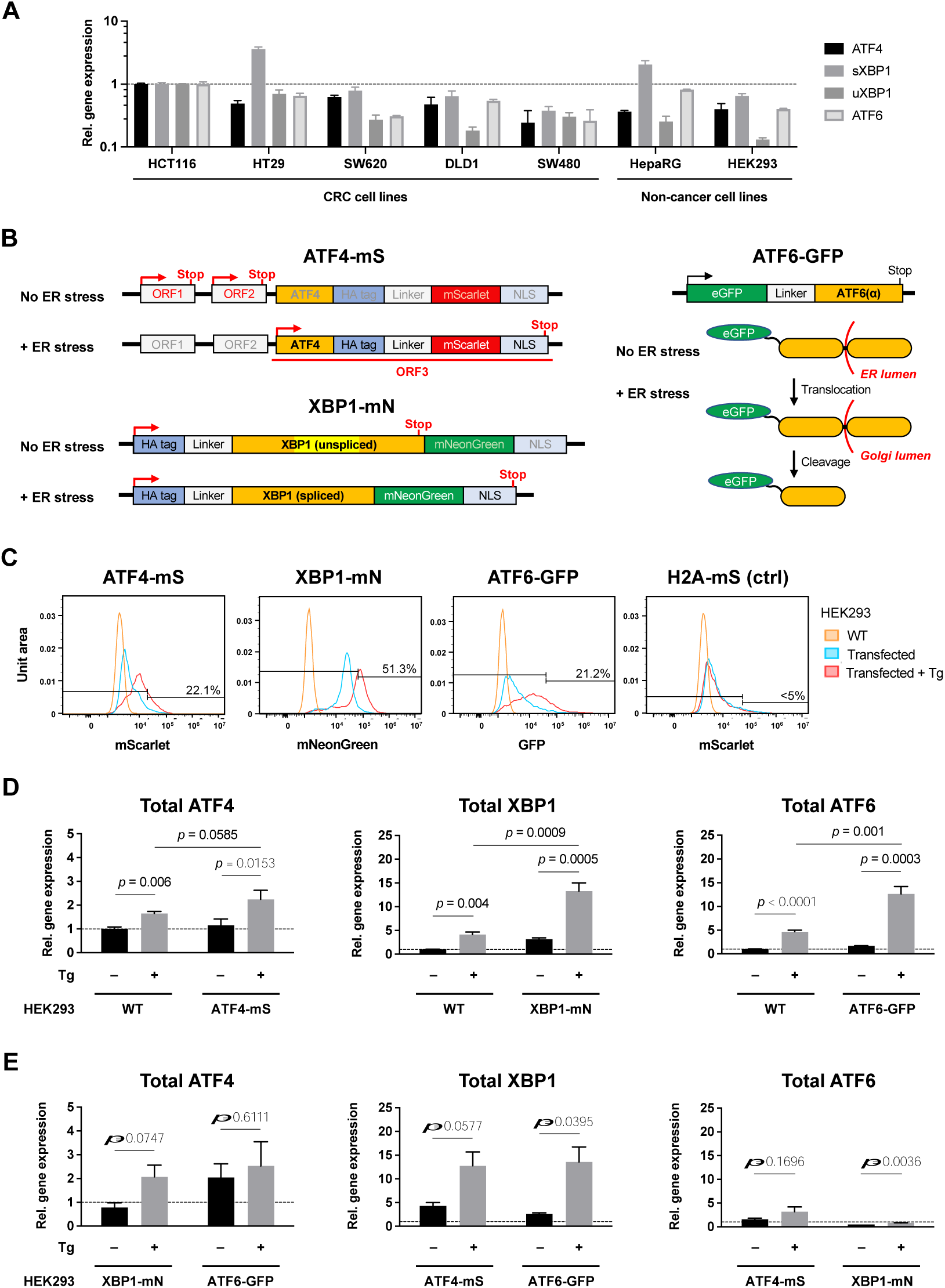
Generation and validation of ER stress reporter cell lines. **(A)** Comparison of basal gene expression levels of ATF4, (s/u)XBP1 and ATF6 between human cell lines. **(B)** Design of ER stress reporter genes encoded with fluorescently labelled ATF4, XBP1 or ATF6. **(C)** The reporter gene expression in transfected HEK293 cells was validated after treatment with Tg (1 μM, 24 h). Cut-off = 95% percentile of transfected cells without Tg treatment. **(D)** Total (endogenous plus exogenous) expression of ATF4, XBP1 and ATF6 in transfected HEK293 cells was analyzed by qRT-PCR after treatment with Tg (1 μM, 24 h). **(E)** Cross effect of the reporter gene expression on endogenous expression of the other ER stress transcription factors. Tg, Thapsigargin.

### Generation and characterization of ER stress reporter cell lines

To monitor and modulate ATF4, XBP1 and ATF6 expression and activation, HEK293 cells, which showed low basal ER stress activity, were stably transfected with plasmid vectors encoding the respective reporter genes that were previously established (Fig. 2B, ATF4-mS, XBP1-mN and ATF6-GFP). Like the endogenous counterparts, ATF4-mS and (s)XBP1-mN are expressed via translational frameshift under ER stress conditions (Nougarède et al., 2018). To mimic the nuclear localization sequence (NLS) found in the native systems, a NLS derived from c-MYC had been added to the C-terminus of ATF4-mS and XBP1-mN fusion proteins, reducing a dispersion of fluorescent signals across the cytosol, as published elsewhere (Walter et al., 2015). ATF6-GFP was shown to undergo the same processing as native ATF6 under ER stress, translocating to the Golgi apparatus, followed by cleavage to release (n)ATF6-GFP to the cytosol, allowing it to then enter the nucleus (Chen et al., 2002). As a control reporter gene unrelated to ER stress signaling, a vector encoding histone H2A fused to mScarlet (H2A-mS) was used. Flow cytometric analysis under canonical ER stress induced by the compound thapsigargin (Tg) confirmed that the reporter genes were expressed and upregulated in transfected cells during ER stress, while the H2A-mS control did not respond (Fig. 2C). Moreover, qRT-PCR confirmed increased expression of total ATF4, XBP1 and ATF6 (endogenous and exogenous) in the transfected cells compared to the wildtype cells (Fig. 2D). Interestingly, there were some cross-branch effects, where expression of a particular reporter influenced the endogenous expression of the other ER stress mediators (Fig. 2E). For instance, the reporter for XBP1 induced overexpression of endogenous ATF4 under thapsigargin-induced ER stress, but not under normal conditions. In contrast, expression of the ATF6 reporter induced endogenous ATF4 gene expression already at basal levels without stimulation. Similarly, the ATF4 and ATF6 reporters both increased endogenous XBP1 gene expression, comparable to the total XBP1 levels detected in the XBP1 reporter line. In contrast, endogenous ATF6 expression levels were not significantly affected by the ATF4 and XBP1 reporters. Thus, the reporter cell lines show robust, yet likely not supra-physiogical levels of the respective ER stress mediators, and can be induced by thapsigargin.

### Modulation of ER stress signaling branches positively affects chemoresistance

Next, the reporter cell lines were assessed for the response to OxaPt to gauge the impact of modulated ER stress signaling branches. XTT assays and the comparison of IC_50_ values showed that the exogenous ER stress reporter gene expression positively affects the resistance to OxaPt (Fig. 3A). Notably, XBP1-mN expression significantly increased OxaPt resistance compared to the wildtype cells, while the H2A-mS control expression showed no significant increase. Changes in total expression of ATF4, XBP1 and ATF6 were evaluated after OxaPt treatment using qRT-PCR (Fig. 3B). In wildtype cells, OxaPt induced significant upregulation of XBP1, while ATF4-mS, XBP1-mN reporter cells did not exhibit upregulation of ATF4, XBP1 or ATF6. ATF6-GFP reporter cells displayed increased expression of all transcription factors in response to the OxaPt-induced stress.

**Fig. 3.**
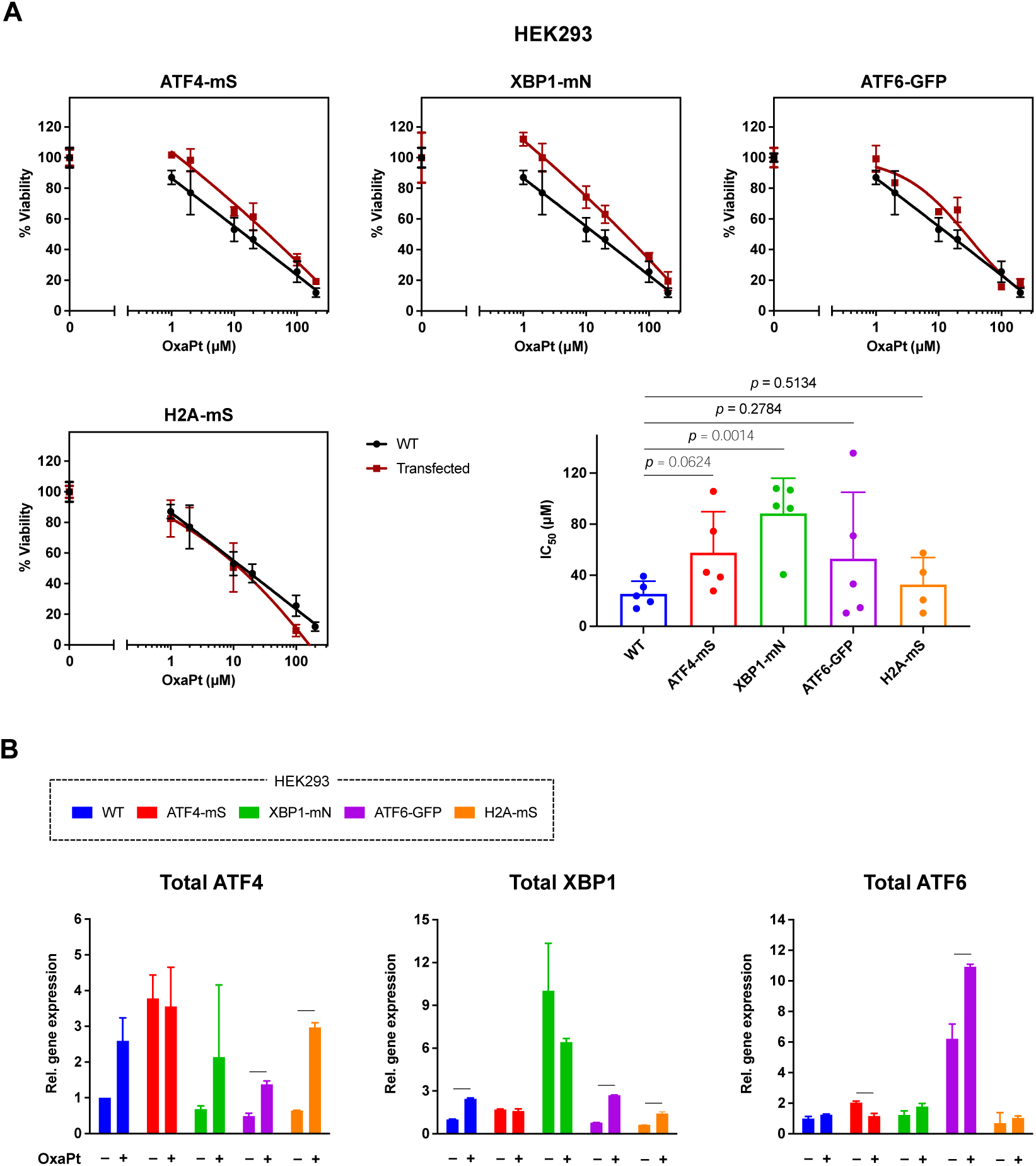
Modulation of ER stress signaling branches positively affects chemoresistance. **(A)** XTT assay of ER stress reporter cells to OxaPt (0–200 μM, 24 h). \**p* < 0.05, \*\**p* < 0.01, compared to HEK293 WT cells. IC_50_ values determined by the XTT assay (*n* = 4– 5) were compared to HEK293 WT cells. **(B)** Total (endogenous plus exogenous) expression of ATF4, XBP1 and ATF6 in HEK293 reporter cells was analyzed by qRT-PCR after treatment with OxaPt (20 μM, 24 h). \**p* ≤ 0.05, ** *p* ≤ 0.01, *** *p* ≤ 0.001, compared to untreated cells. OxaPt, Oxaliplatin.

### Inhibition of ATF6 sensitizes ER-stressed cells to chemotherapy

Parental HCT116 and HEK293 cells were tested for pharmacological inhibition targeting the three ER stress signaling branches (GSK2656157, STF-083010 and Ceapin-A7) to assess the impact on OxaPt resistance. GSK2656157 selectively deactivates PERK phosphorylation, interfering with the p-eIF2α/ATF4 pathway (Axten et al., 2013). STF-083010 (STF’010) inhibits IRE1α endonuclease, preventing mRNA splicing pf XBP1, but without affecting its kinase activity (Papandreou et al., 2011). Ceapin-A7 is known as an ATF6 inhibitor, specifically blocking the ER-to-Golgi transport of ATF6 (Gallagher et al., 2016). Cell viability after treatment with inhibitors alone or in combination with OxaPt was assessed using XTT proliferation assays (Fig. 4A). In HCT116 cells with high basal ER stress activity (Fig. 1A, B), ATF6 inhibition increased the sensitivity against OxaPt. In contratst, inhibition of ATF4 and XBP1 had the opposite effect and further increased OxaPt resistance (Fig. 4A, HCT116). In HEK293 cells with lower basal ER stress activity (Fig. 1A, B), treatment with either of the ER stress inhibitors led to increased resistance against OxaPt (Fig. 4A, HEK293). In accordance, flow cytometric analysis demonstrated an increase in apoptosis, detected as AnnexinV-positive cells, in OxaPt treated HCT116 cells upon combination with ATF6 inhibition (Fig. 4B). The specific effect of pharmacological ATF6 inhibition on OxaPt resistance was further confirmed in human colon organoids using cytotoxicity assays based on SYTOX staining, a DNA dye that exclusively penetrates damaged cell membranes (Fig. 4C; Fig. S3).

**Fig. 4.**
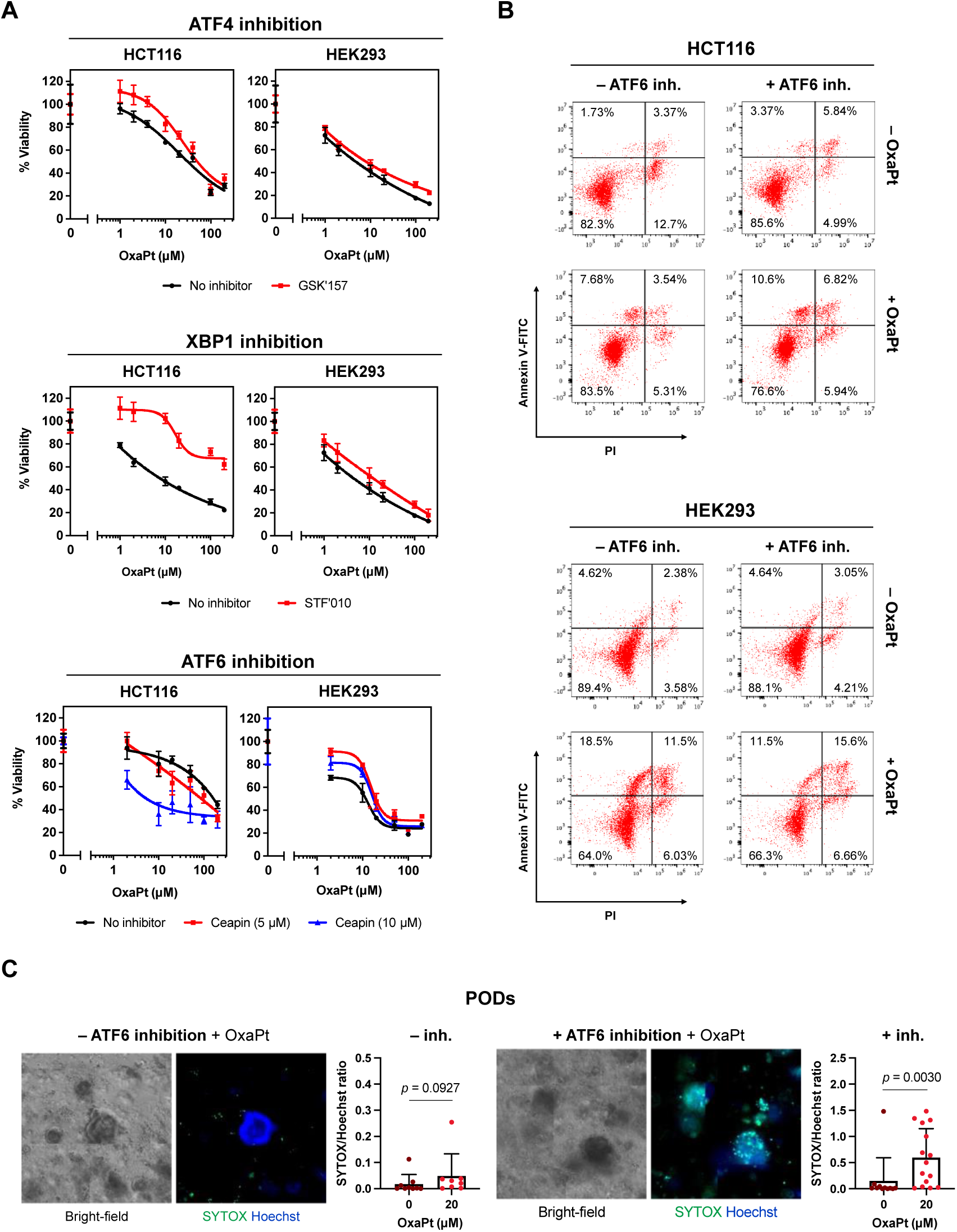
Inhibition of ATF6 sensitizes ER-stressed cells to chemotherapy. **(A)** XTT cell viability assay of HCT116 and HEK293 cells treated with inhibitors for ER-stress receptors and OxaPt. HCT116 and HEK293 cells were treated with GSK2656157 (GSK’157, 10 μM, 1 h), STF083010 (STF’010, 50 μM, 1 h) or Ceapin-A7 (Ceapin, 5 or 10 μM, 6 h) prior to the treatment with OxaPt (0–200 μM, 24 h). *p ≤ 0.05, **p ≤ 0.01, ***p ≤ 0.001, \*\*\*\**p* ≤ 0.0001. **(B)** Flow cytometric analysis of apoptosis in HCT116 and HEK293 cells treated with Ceapin-A7 (ATF6 inh., 5 μM, 1 h) prior to the treatment with OxaPt (20 μM, 24 h). **(C)** Patient-derived organoids (PODs) were treated with Ceapin-A7 (5 μM, 6 h) prior to the treatment with OxaPt (0–20 μM, 48 h), followed by co-staining with SYTOX and Hoechst. SYTOX/Hoechst ratios (each point represent one organoid) were compared. Organoids were isolated from adjacent normal colon tissues from a stage IV CRC patient. * *p* ≤ 0.05, ** *p* ≤ 0.01, *** *p* ≤ 0.001, compared to the vehicle control (DMSO). OxaPt, Oxaliplatin.

### Oxaliplatin induces a rapid ER stress response that differs from canonical induction by thapsigargin

We analyzed kinetics of the ER stress signaling activation in live HEK293 cells under OxaPt treatment, compared to the canonical ER stress responses induced by thapsigargin (Tg) (Fig. 5). Western blotting revealed that OxaPt quickly triggered cellular responses, as characterized by cleavage of poly(ADP-ribose) polymerase 1 (PARP1), a typical marker of DNA damage repair pathways and apoptosis. Unlike Tg-induced canonical proteostatic responses, OxaPt did not lead to upregulation of the ER-mediated chaperone BiP at protein level (Fig. 5A). PARP1 cleavage coincided with ATF6 cleavage, while eIF2α phosphorylation lagged behind these events. The splicing assay for XBP1 mRNA indicated that XBP1 activation also occurred later than ATF6 cleavage under conditions of OxaPt-induced ER stress (Fig. 5B). Immunofluorescence microscopy allowed a detailed analysis of ATF6-GFP reporter protein subcellular distribution. Using counterstaining with DNA dyes (DAPI) and a Golgi apparatus marker (Golgin-97), we segmented and quantified the fluorescence signal of ATF6-GFP in individual cells at defined treatment time points (Fig. S4). Subcellular colocalization analysis of ATF6-GFP reporter cells revealed rapid translocation of ATF6-GFP to the Golgi apparatus during OxaPt-induced ER stress compared to Tg-induced stress. In contrast, no time-dependent changes were observed in H2A-mS control cells during both treatments (Fig. 5C–D). Of note, morphological changes characterized by reduced roundness of nuclei and nuclear shrinkage were observed after 6 hours of OxaPt treatment, which were not evident in Tg treatment (Fig. 5C).

**Fig. 5.**
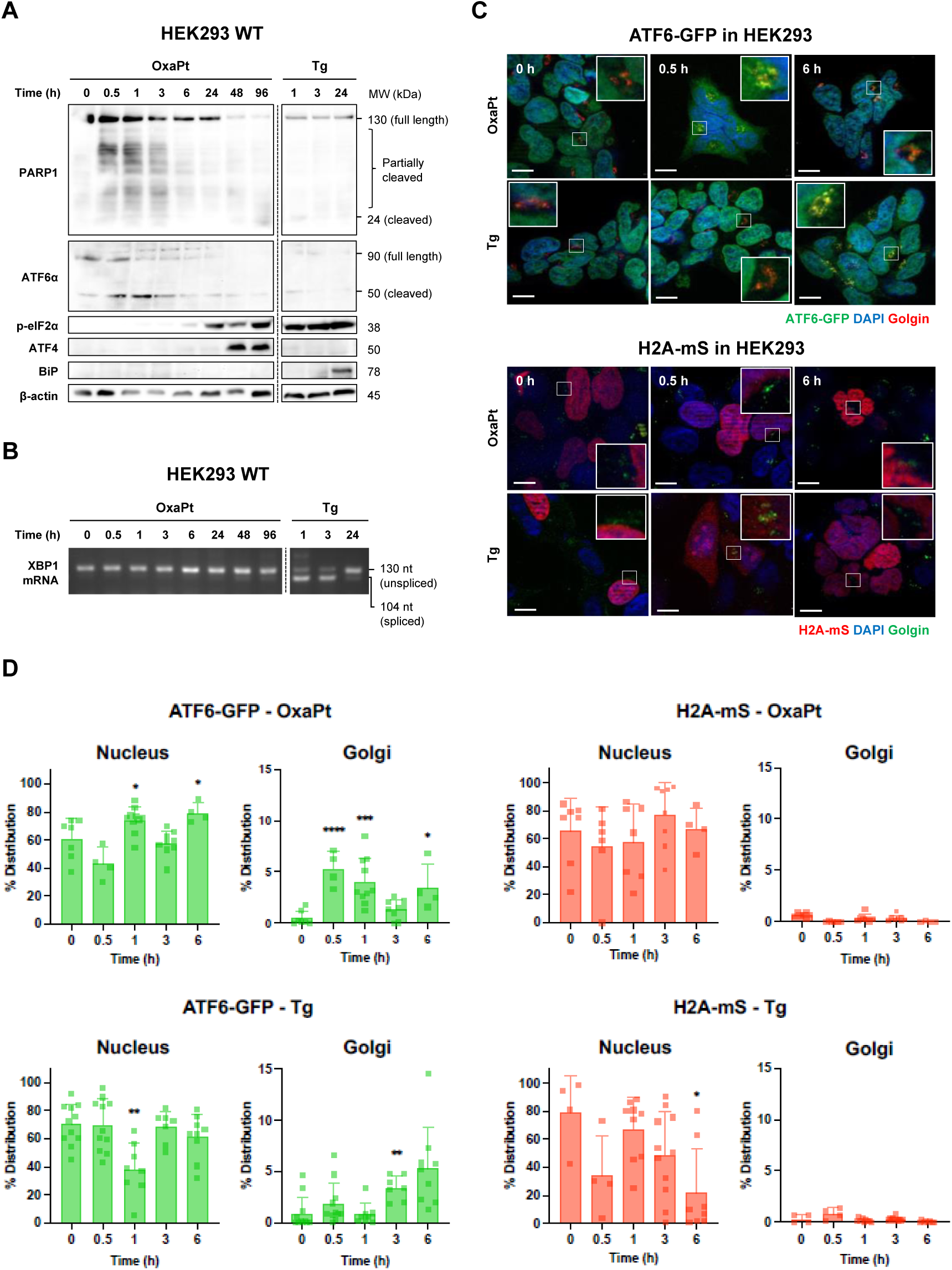
Chemotherapy rapidly induces the ER stress signaling in non-canonical way. Time-lapse analysis of the ER stress inducible markers in HEK293 cells treated with OxaPt (20 μM, 0–96 h) or Tg (1 μM, 0–24 h) by **(A)** Western blotting and **(B)** XBP1 mRNA splicing assay. **(C)** Optical section images of HEK293 ATF6-GFP reporter and H2A-mS control cells after treatment with OxaPt (20 μM, 0–6 h) or Tg (1 μM, 0–6 h). The cells were co-stained with DAPI and anti-Golgin-97 antibody. Scale bar = 10 nm. **(D)** Quantification of ATF6-GFP reporter localization to nucleus or Golgi regions in HEK293 cells was evaluated and compared that of H2A-mS reporter in respective cells based on the optical section images (workflows are depicted in Fig. S5). \**p* ≤ 0.05, \*\**p* ≤ 0.01, \*\*\**p* ≤ 0.001, \*\*\*\**p* ≤ 0.0001, compared to the 0 h treatment. OxaPt, Oxaliplatin; Tg, Thapsigargin.

### Proteome-wide analysis shows that response to chemotherapy differs between the ER stress signaling branches

To further characterize the global cellular impact of chemotherapy and ER stress reporter gene expression, we conducted a proteome-based analysis using parental HEK293 wildtype cells, XBP1-mN and ATF6-GFP reporter cells, since these two lines showed the robust effects on chemosensitivity. Mass spectrometry based analysis revealed significant down/upregulation in response to OxaPt treatment, with some overlap between the groups (Fig. 6A). We also identified proteins showing significant changes upon Tg treatment for each cell line (Fig. S5A), partially overlapping with those in the OxaPt-treated cells (Fig. 6B). For comparison, proteins exhibiting significant changes during massive apoptosis induced by staurosporine in wildtype cells were also identified, and the overlap with those affected by OxaPt and Tg treatments was assessed (Fig. S5B; Fig. S6C, left panels). We compared the proteome of each cell line treated with OxaPt, focusing on functional enrichment of downregulated and upregulated pathways. This analysis revealed distinct responses among cell lines: wildtype cells displayed downregulation of functions related to the nucleolus, microtubules, and ribosome biogenesis (Fig. S6A, right panel). XBP1-mN cells exhibited upregulation in lysosome machinery and amino acid metabolism, whereas ATF6-GFP cells displayed a different set of upregulated pathways, including DNA damage responses (Fig. 6C–D, right panels). Of note, Tg induced similar proteostatic responses in these cell lines, and wildtype cells showed minimal changes in the global proteome even with a relaxed significance threshold, *q* ≤ 0.2 (Fig. S6B; Fig. S7A–B, right panels).

**Fig. 6.**
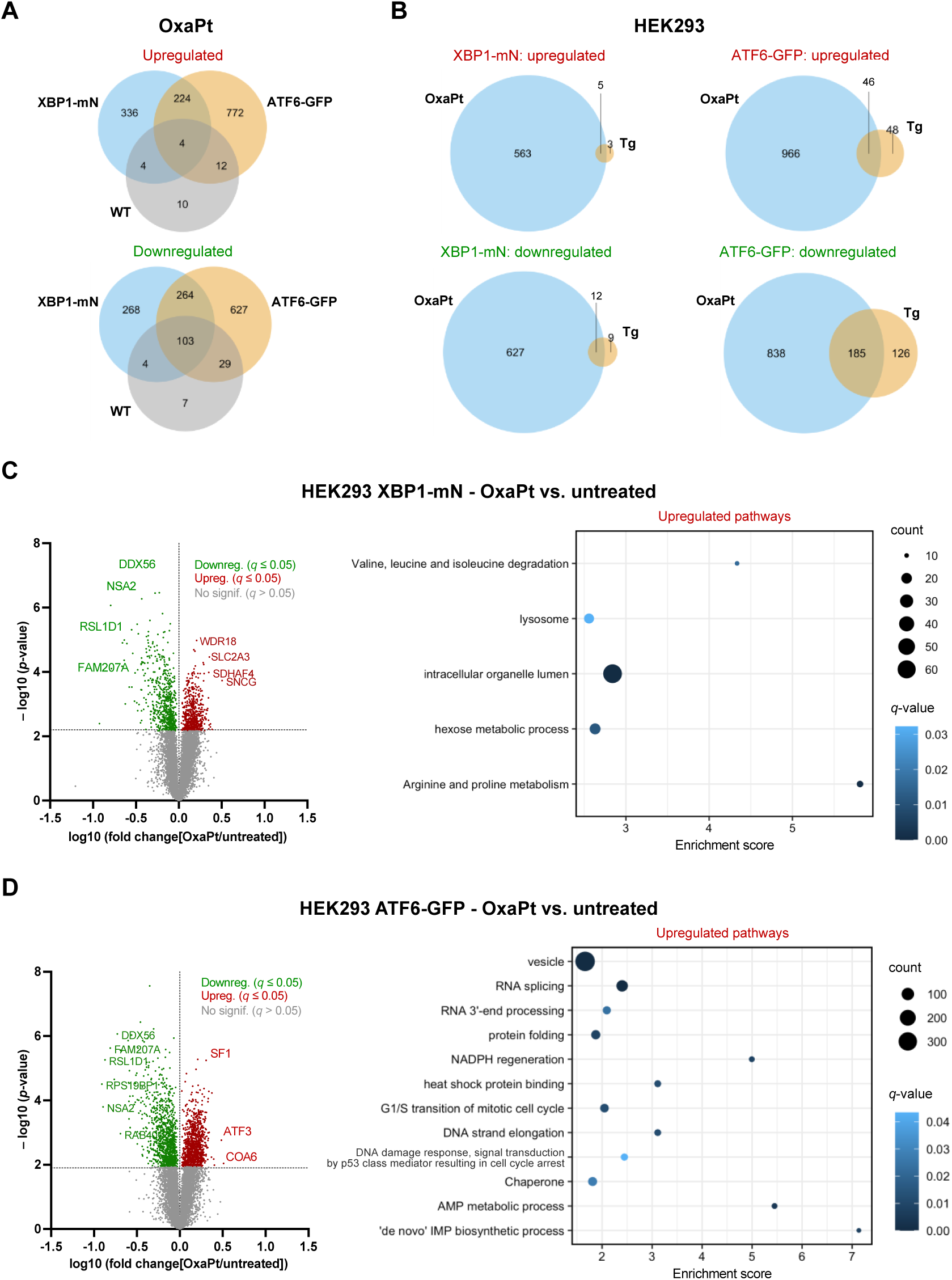
The XBP1- and ATF6-dependent modulation induces different downstream signaling effect on Oxaliplatin resistance. **(A)** The numbers of significantly down/upregulated proteins (*q* ≤ 0.05) in HEK293 WT, XBP1-mN and ATF6-GFP reporter cells treated with OxaPt (20 μM, 24 h), compared to untreated condition, and overlaps of those for each cell line. **(B)** The numbers of significantly down/upregulated proteins (*q* ≤ 0.05) in XBP1-mN and ATF6-GFP reporter cells treated with OxaPt (20 μM, 24 h) or Tg (1 μM, 24 h), compared to untreated condition, and overlaps of those for each treatment within each cell line. **(C, D)** Volcano plots of down/upregulated proteins in (C) XBP1-mN or (D) ATF6-GFP cells, compared to untreated conditions (left panels) and enrichment pathway analysis based on gene ontology terms performed for upregulated proteins (right panels, *q* ≤ 0.05). OxaPt, Oxaliplatin.

### The expression of ATF6 reporter increases radiotherapy resistance

The results above collectively suggest a putative association between the ATF6 level and cellular DNA damage repair following chemotherapy *in vitro*. Therefore, the effect of the ER stress reporter expression on cellular response to radiotherapy was investigated in detail since radiation is known to induce DNA damage, particularly double-strand breaks (DSB), which are considered the most lethal form of DNA damage to cells (Helleday et al., 2008; Valerie & Povirk, 2003). Control cells and ATF6-reporter cells were subjected to specific doses of irradiation (0–8 Gy). qRT-PCR analysis revealed that total ATF6 expression was notably higher compared to wildtype cells and increased in a time-dependent manner, while ATF4 and XBP1 expression remained comparable to wildtype cells and showed minimal response to radiation treatment (Fig. 7A). Flow cytometry following X-ray irradiation demonstrated a dose-dependent induction of ATF6-GFP reporter expression (Fig. 7B). Annexin-PI flow cytometric assay indicated that radiation induced apoptosis in a dose-dependent manner (Fig. 7C). ATF6-GFP cells and H2A-mS control cells exhibited significantly more robust colony formation after irradiation compared to wildtype cells (Fig. 7D, *p* = 0.0240 and 0.0173, respectively).

**Fig. 7.**
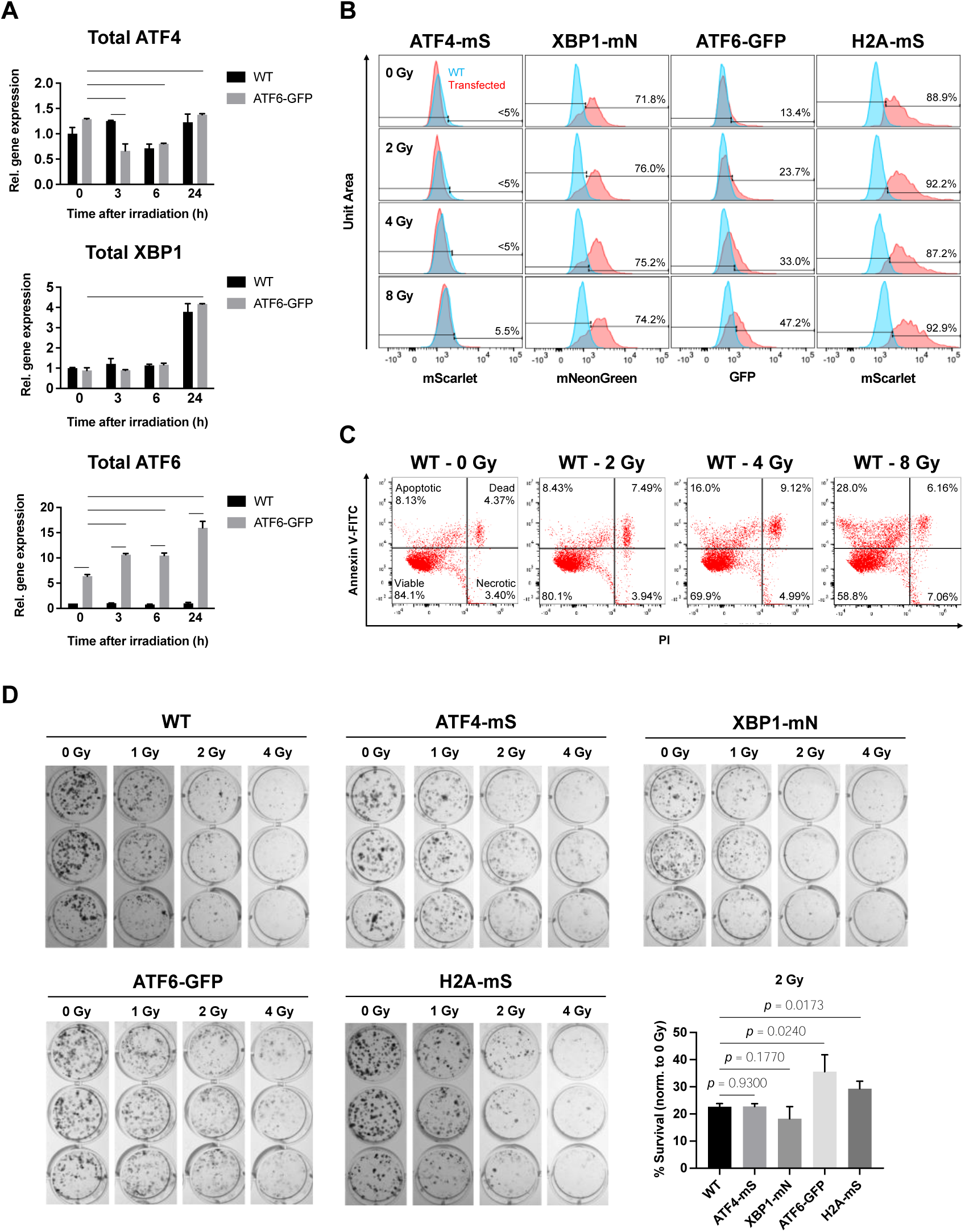
The expression of ATF6-GFP reporter increases radioresistance. **(A)** qRT-PCR analysis of the total (endogenous plus exogenous) expression of ATF4, XBP1 and ATF6 in HEK293 reporter cells 24 h after irradiation at 2 Gy. \**p* ≤ 0.05, \*\**p* ≤ 0.01, \*\*\**p* ≤ 0.001, compared to that of 0 h after treatment. **(B)** Flow cytometric detection of the reporter gene expression in HEK293 reporter cells 30 h after irradiation at 0–8 Gy. **(C)** Flow cytometric analysis of apoptosis in HEK293 cells 48 h after irradiation at 0–8 Gy. **(D)** Colony formation assay of HEK293 reporter cells irradiated at 0–4 Gy and comparison of the survival rates between the cells irradiated at 2 Gy.

## Discussion

Predicting responses to cancer treatment, such as chemo- and radiotherapy, is challenging due to stress-resistant tumor subpopulations selected through adaptive responses, for which molecular mechanisms and clinical impacts are still largely unclear (Holohan et al., 2013). The ER stress response is among these adaptive processes. Since it was initially characterized as a compensatory mechanism for proteostasis (Kaufman, 2002; Mori, 2000), many studies have suggested its active involvement in other cellular processes, constituting a central hub that integrates vital pathways for cancer cells to survive a hostile microenvironment and intracellular oncogenic changes (Dicks et al., 2015; González-Quiroz et al., 2020). Indeed, activation of the ER stress signaling has been observed in many types of human tumors (Wang & Kaufman, 2014) and is associated with highly proliferative, malignant and aggressive properties in cellular and animal models (Urra et al., 2016). However, due to a lack of systematic evidence on which signaling branches are essential for therapy resistance, the underlying mechanisms remain unclear. Among the various approaches to monitor ER stress in live cells, the use genetic reporter systems is well established and has the advantage to allow the study of cellular reactions under real-time conditions and in vivo (Sicari et al., A guide to assessing endoplasmic reticulum homeostasis and stress in mammalian systems. The FEBS Journal 287 (2020) 27–42). Of course, the stable introduction of a fluorophore-coupled, yet fully functional ER stress mediator, may lead to its expression above physiological levels. However, our analysis shows that this does not seem to be the case, when assessing mRNA expression of endogenous and exogenous ER stress mediators (Fig. x), as well as global proteome results which show essentially no changes in otherwise untreated ER stress reporter cell lines, compared to the parental cell line.

In this study, we explored the individual roles of ER stress signaling branches in chemo- and radiotherapy resistance of CRC, emphasizing the nuclear activity of key transcription factors ATF4, sXBP1 and nATF6. The analysis of cancer patients in TCGA dataset clearly showed that ATF6 is predominantly amplified at a genome level in several prevalent types of solid cancer. Genomic amplifications, especially of ATF6, were much more frequently observed than other genetic changes (*e.g.*, point mutations, genomic losses), hinting towards a “gain of function” type of alteration for ATF6. Of note ATF6 was amplified in 36% of all cases in a pan-cancer cohort. In colorectal cancer, 23% of cases showed a gene-level amplification, which is comparable to an earlier observation that a subgroup of CRC patients present with ATF6 gene alterations with negative prognostic association (Coleman et al., 2018). Of course, genome level alterations do not necessarily correspond to changes on transcriptional or translational level, nor the cellular activity. Interestingly, whereas all three ER stress mediators were negatively associated with overall survival, ATF6 gene amplification was unique in its correlation with worse disease-free survival, underscoring the putative role of ER stress, and specifically ATF6, in the context of disease outcome and therapy responses.

Furthermore, positive ATF6 was shown connected with increased genomic instability both in genetic and protein levels. TCGA analysis of amplified ATF6 concurred with TP53 mutation in association with aneuploidy and prognosis indicated that amplified ATF6 is related to poor patient outcome, particularly in the context of TP53-driven chromosomal instability. ONLY IN PAN-CANCER?

Additionally, flow cytometric analysis of a small cohort of surgically resected human CRC showed that intratumoral ATF6 protein expression levels were increased in MSI-high cases, while ATF6 levels in adjacent normal colon remained unchanged, as well as XBP1. Genetic instability comes in various forms, in CRC as chromosomal instability (CIN) with frequent aneuploidy, and as microsatellite instability in a subset of cases. Due to the limited sample size in the present study, no definitive conclusions can be drawn, but it appears as if high levels of ATF6 are specifically associated with increased genetic instability in CRC, and it is safe to assume that tumor cell clones with aneuploidy gain survival advantages upon acquisition of gene-level gains of ATF6.

MSI can be translated into an accumulation of DNA replication errors and a high frequency of frameshift mutations, causing cell-intrinsic genotoxic stresses (Jiricny, 2006). In fact, the number of MSI in specified sites, has been regarded as a predictive marker for the response to 5-fluorouracil (5-FU)-based adjuvant chemotherapy for stage II-III CRC (Battaglin et al., 2018; Tomasello et al., 2022; Tougeron et al., 2016). MSI-high cases are generally much more frequent in right-sided CRC tumors such as in coecum and colon ascendens. However, the MSI cases in this study were all left-sided (*n* = 3). Therefore, it should be noted that the MSI dataset may be biased and not represent MSI in the whole CRC population. Collectively, these clinical data raise the hypothesis that tumors that harbor genomic instability can drive selective pressure towards amplification of the ATF6 gene locus, thereby promoting cell survival in highly stressful conditions. In accordance with this model, recurrent tumors exhibited high ATF6 expression levels (Ginos et al., 2004), and ATF6 levels were found to be correlated with poor prognosis in CRC (Lin et al., 2007). Previous studies have also suggested that the activation of ER stress signaling interacts with DNA damage response (DDR) systems and may suppress the onset of genomic instability (Bobrovnikova-Marjon et al., 2010; Henry et al., 2010). However, regarding the role of ATF6, our current understanding is very limited, despite the recent report that ATF6 can sustain the expression levels of DNA repair proteins by activating the mammalian target of rapamycin (mTOR) signaling, which can protect colon cancer cells from the cytotoxic effect of ER stress-targeting drugs such as Tg (Benedetti et al., 2022). Therefore, the findings here might represent a novel and clinically relevant connection between the ATF6 branch and genomic instability.

In addition to the clinical data, cellular *in vitro* models, as well as primary *ex vivo* organoid models, also highlighted a putative connection between the activation of ER stress transcription factors and responses to chemo-/radiotherapy. We established stable reporter cell-line systems to monitor and modulate the activation of ER stress proteins in living cells in response to cancer therapies. Indeed, the reporter lines showed specific ER stress reporter induction upon canonical proteotoxic stress with thapsigargin and significantly increased translocation of ATF6 to the Golgi apparatus and nucleus. Notably, the data on the reporter cell lines gathered here collectively showed that the endogenous as well as the exogenously expressed proteins behave quite similarly in terms of gene/protein expression, subcellular localization and downstream effects. Thus, we attempted to compare the kinetics of ER stress induction between different treatments and screened for the cellular responses specific to the individual signaling branches. A fluorescent probe attached to a protein of interest was utilized as approach, under control of the tomegalovirus (CMV) promoter, which generally drives a strong increase in the amount of a protein (Qin et al., 2010). Therefore, the models generated here might present an expression of the ER stress markers at supra-physiological levels. However, previously, a subset of human CRC patients with decreased prognosis was found with an up to ten-fold increase of nATF6 on protein level in primary colorectal cancer tissue, indicating that indeed, a strong gain of expression of ER stress markers may be also observed in clinical samples (Coleman et al,. 2018). Therefore, we employed these reporter cell lines as “Gain of function” models for these transcription factors. The reporter gene expression positively affected resistance to OxaPt, among which the XBP1-mN reporter cells showed a significant increase of resistance to OxaPt compared to the wildtype.

On the other hand, ATF6-GFP reporter cells showed a significant increase in survival from radiotherapy, in which DDR is emphasized. Of note, the H2A-mS control cells, which exogenously expressed histone H2A independently of ER stress, did not show many ER-stress specific effects on chemotherapy resistance, indicating that exogenous protein overexpression and likely the resulting additional proteotoxic stresses are negligible in this experimental system. Therefore, these reporter cell lines work as overexpression models for the ER-stress transcription factors, which revealed that XBP1-dependent modulation actively promoted chemoresistance, while ATF6-dependent modulation increased radiotherapy resistance. According to the proteome-based analysis of downstream pathway enrichment, the cellular responses to OxaPt were different from the canonical response to proteotoxic stress with Tg and also different between the reporter cell lines: the ATF6-dependent modulation appeared to be involved in DNA damage repair, while XBP1 branches are relatively multi-functional and involved in ribosomal biogenesis stress. These results correspond to immunoblot analysis of HEK293 wildtype cells, in which ATF6 is more closely involved in PARP1 cleavage-induced cellular responses than ATF4 and XBP1. PARP1 cleavage is a hallmark of p53-dependent DDR (Amé et al., 2004). Reportedly, while HEK293 cells constitutively express some viral oncoproteins that compromise function of p53-mediated DNA damage repair, conferring insensitivity to structural and numerical CIN (Bester et al., 2011; Komorek et al., 2010), expression of p53-regulated genes in response to DNA-damaging agents has been demonstrated (Jeon et al., 2012; Kishore et al., 2007). Therefore, the immunoblot data supports the proteome-based results that ATF6 can activate DDR in response to OxaPt treatment.

The results above suggest that ATF6 is responsible for early DDR, while the XBP1 branch is rather multi-functional and involved in ribosomal biogenesis stress during chemotherapeutic treatment with OxaPt. Ribosomal biogenesis is a very tightly regulated process for protein translation, which involves the production of ribosomal proteins and rRNA encoded by RNA polymerases (Pol). The first step of rDNA transcription induced by RNA Pol I leads to the production of the 47S precursor rRNA (pre-rRNA), which will then be cleaved and assembled with ribosomal proteins to form the 40S and 60S subunits. This step, occurring within sub-compartments of the nucleolus, is affected by the intracellular nutrimental conditions and the activity of oncoproteins such as c-Myc and p53, triggering nucleolar stress leading to cell cycle arrest and/or apoptosis (Russo & Russo, 2017). Hyperactivation of rRNA processing and ribosomal biogenesis plasticity are hallmarks of malignant tumor progression that is essential for increased protein synthesis to maintain unremitting cancer cell growth, emerging as novel therapy targets for human CRC (Drygin et al., 2010; Ferreira et al., 2020).

To the best of our knowledge, the roles of the ER stress signaling branches in these processes and the mechanisms for the acquisition of resistance still need to be explored except for the PERK-ATF4 axis or its proximal pathways. One study showed that PERK-mediated ER stress signaling promoted a p53-dependent cell-cycle arrest by affecting interaction between several ribosomal proteins (Zhang et al., 2006). Another related study is on characterizing the general control nonderepressible 2 (GCN2), a cytosolic kinase that senses a lack of amino acids and converges on the phosphorylation of eIF2α, in which GCN2 was shown to differentially regulate ribosomal biogenesis in colon cancer cells depending on the nutrient availability. Furthermore, pharmacological co-inhibition of GCN2 branches and RNA polymerase I activity enhanced the effect of chemotherapy including OxaPt on patient-derived colon tumor organoids (Piecyk et al., 2023). The present work highlighted the roles of XBP1 and ATF6 in the ribosomal biogenesis stress. Through the microscopy of HEK293 cells treated with OxaPt, morphological changes of the cell nuclei were observed that are typical for damages from nucleolar stress (Yang et al., 2018). Moreover, the findings that the XBP1-dependent modulation induced a more pronounced increase in the resistance to OxaPt than the ATF6-dependent modulation also suggest that induction of ribosomal biogenesis stress by OxaPt is more emphasized than the DNA damage responses when treated for a longer time. Therefore, it may be concluded that activation of the XBP1-dependent signaling pathways is a dominant factor in the resistance to OxaPt. However, pharmacological inhibition of the IRE1-XBP1 signaling pathways displayed no apparent decrease in the chemoresistance or even a reverse effect, although inhibition of the ATF6 signaling pathways sensitized CRC cells to OxaPt, which were recapitulated on human colon organoids independently. One possible explanation is that XBP1 plays crucial roles in ribosomal biogenesis stress response but converges on multiple shared downstream effector molecules, and therefore, inhibition of XBP1 can be compensated by other signaling pathways, while inhibition of ATF6 has much impact because it plays non-redundant key roles in the DDR. Another explanation is based on the assumption of a time lag between the acute DDR and the subsequent ribosomal biogenesis stress responses, which was observed in the immunoblot analysis. Cells first activate DDR for adaptation to the genotoxic aspect of OxaPt, and ribosomal biogenesis stress manifests later on. Thus, inhibition of the ATF6 branch has an immediate impact on the cytotoxic effect of OxaPt. On the other hand, inhibition of IRE1-XBP1 does not much affect the OxaPt-induced ribosomal biogenesis stress responses. It might be because adaptation to the initial genotoxic stress in an ATF6-dependent manner renders the cells less dependent on XBP1 function, activating compensatory pathways. On the other hand, the mTOR pathway also reciprocally controls the RNA Pol I activity (Chen et al., 2016). Such an indirect connection between ATF6 and ribosomal biogenesis implicates the existence of compensatory pathways for ribosomal biogenesis stress adaptation independent of XBP1 branches. Of note, the activity of IRE1 inhibitor has been validated on several other human cell lines in previous research, with essentially similar concentrations and time scales (e.g., 60 μM, 18 h; (Papandreou et al., 2011)), and therefore, it is unlikely that the inhibitor loses its activity before the ribosomal biogenesis stress arises. In a context in which genotoxic stress induction is more dominant, such as DSBs induced by radiotherapy, the roles of ATF6 in DDR can be more emphasized. Collectively, the contribution of individual ER stress signaling branches is highly dependent on the cancer therapies, which were explained through potentially dominant mechanisms for each treatment.

OxaPt has a broad spectrum of impact on cellular adaptive signaling, including DDR and ribosomal biogenesis stress responses. In association with that, OxaPt-based chemo(radio)therapy harbors a risk for developing a multi-drug resistance, which shows cross-resistance to many other chemically and functionally related drugs and inadvertently reduces the options for curative and palliative treatment (Hsu et al., 2018). Therefore, it might be beneficial to stratify cancer patients who potentially show poor response to OxaPt and can eventually develop resistance, aside from the other patients, before or early on in treatment, and to consider other therapeutic strategies, such as targeting ribosomal biogenesis in cancer cells, which has gained considerable attention as a novel approach in the treatment of human CRC. For example, CX-5461 and CX-3543, new drugs that selectively bind to rDNA-enriched G-quadruplex, have shown more cytotoxicity in particular cancer cells that have high activity of ribosomal biogenesis, compared to existing chemotherapy such as OxaPt that can also affect rRNA processing in an unselective manner (Drygin et al., 2010; Ferreira et al., 2020). CX-5461 is in phase I/II clinical trials for hematological cancers (Khot et al., 2019), but further mechanistic studies are necessary because, of interest, it has also shown *in vitro* and *in vivo* that cytotoxicity of CX-5461 and CX-3543 is mediated by the activation of a robust DDR, indicating crosstalk between ribosomal biogenesis inhibition and DDR activation (Xu et al., 2017).

Two hypothetical models were thus raised, based on the clinical and CRC cell-based analysis: cellular responses to the initial genotoxic stress of OxaPt occurs in an ATF6-dependent manner, and it reprograms the adaptive signaling pathways for the subsequent ribosomal biogenesis stress into less dependence on XBP1 function.

Of note, this effect was significantly observed in HCT116 cells, whose basal activity of the ER stress signaling is relatively high, but not in HEK293 cells, which showed less basal activation of the ER stress, indicating that ER stress-targeting treatment is more effective for cancer cells which adapted to the ER stress. It collectively suggests that tumors which developed genomic instability can show a positive response to OxaPt-induced cytotoxicity in ATF6 inhibition rather than in XBP1 inhibition, and therefore, it will be meaningful to further investigate whether ATF6 inhibition potentiates ribosomal biogenesis-targeting therapy.

In conclusion, the activation status of the ER stress signaling branches in tumor represents the therapy response in association with genomic instability, and therefore, is promising as a predictive biomarker for the selection of better treatment options for individual patients. Furthermore, intervention with them can modulate the cytotoxicity of OxaPt treatment, and the mechanistic insight into the OxaPt-induced cytotoxicity opened a strategic window for ATF6 inhibition to combine with ribosomal biogenesis-targeting therapy.

---

**Table S1.** Correlation between XBP1 or ATF6 positivity and clinical characteristics in cohort 1 (*n* = 7): human CRC tumors and adjacent normal colon tissues (Mann-Whitney test).

|  |  |  | Comparison between two groups - *p*-values |  |  |  |
| --- | --- | --- | --- | --- | --- | --- |
|  |  |  | XBP1 |  | ATF6 |  |
|  |  |  | Normal | Tumor | Normal | Tumor |
| Clinical characteristics | All | MSI |  |  |  |  |
| Gender - no. of patients |  |  |  |  |  |  |
| Male | 3(2) | 1 | 0.5147 | 0.8508 | 0.2137 | 0.5930 |
| Female | 4 | 2 |  |  |  |  |
| Age - years |  |  |  |  |  |  |
| Median | 73 | 66 | — | — | — | — |
| Interquartile range | 66–80 | 66–74 |  |  |  |  |
| Location - no. of patients |  |  |  |  |  |  |
| Right-sided | 2(1) | 0 | 0.5232 | 0.8918 | 0.7562 | 0.3726 |
| Left-sided | 5 | 3 |  |  |  |  |
| MS status - no. of patients |  |  |  |  |  |  |
| MSS | 4(3) | — | 0.8460 | 0.5532 | 0.5211 | *0.0195 |
| MSI | 3 | — |  |  |  |  |
| UICC stages - no. of patients |  |  |  |  |  |  |
| I | 2(1) | 1 | — | — | — | — |
| II | 2 | 0 |  |  |  |  |
| III | 2 | 1 |  |  |  |  |
| IV | 1 | 1 |  |  |  |  |
| pT stages - no. of patients |  |  |  |  |  |  |

TNM: tumor lymph node metastasis; (): the sample number for the ATF4 and XBP1 datasets.

