## Supplementary Figure Legends for "Endoplasmic reticulum stress signaling actively contributes to therapy resistance in colorectal cancer"

**Supplementary Data**

**Supplementary Figure Legends**

**Fig. S1 Association of ATF4, XBP1 and ATF6 positivity with clinical characteristics** **derived from TCGA datasets and surgically resected human CRC.** **(A)** Kaplan-Meier analysis of disease-free survival (DFS) among cancer patients with upregulated all three genes (either ATF4, XBP1 or ATF6) (left top) and overall survival (OS) among cancer patients with upregulated ATF4, XBP1 or ATF6, compared to patients with no alterations from the public TCGA datasets. **(B, C)** Positivity of ATF4, XBP1 and ATF6 was assessed by flow cytometry and compared between matched pairs of colon tumor and adjacent normal tissues (*n* = 3). **B** MSI group; **C** MSS group. **(D)** Comparison between colon tumors at T1–2 stages and T3–4 stages. **p* ≤ 0.05, compared to the control group. MSI, microsatellite instable; MSS, microsatellite stable or below the detection limit.

**Fig. S2** **Connection between basal activity of the ER stress signaling and chemoresistance of human cell lines**. **(A)** Western blotting of subcellular fractions from HCT116, HT29 and DLD1 cells. WCL, whole cell lysate; Nuc, nucleus fraction; Cyto, cytosolic fraction. **(B)** MTT assay of 7 human cell lines to OxaPt (0–2 mM, 24 h). **(C)** Correlation between the basal gene expression of ATF4, (s/u)XBP1, ATF6 and IC_50_ values determined by MTT assays of 6 human cell lines to OxaPt (0–2 mM, 24 h). r, Pearson’s correlation coefficient. CRC, colorectal cancer; OxaPt, Oxaliplatin.

**Fig. S3** **Effect of pharmacological inhibition of ER stress signaling branches on chemotherapy responses in cells and human colon organoids.** SYTOX-Hoechst staining of human colon organoids pretreated with GSK2656157 (10 μM, 1 h), STF083010 (50 μM, 1 h) and Tg (1 μM, 1 h), followed by treatment with OxaPt (0, 20, 200 μM, 48 h). Organoids were isolated from adjacent normal colon tissues from a stage I CRC patient. ** *p* ≤ 0.01, *** *p* ≤ 0.001, **** *p* ≤ 0.0001, compared to the vehicle control (DMSO). OxaPt, Oxaliplatin.

**Fig. S4 The workflow for the analysis of co-localization of reporter proteins with nucleus and Golgi regions based on optical section images.** The ROIs for individual cells were manually assigned based on bright-field images, and co-localization of signals from two reporter channels was quantified.

**Fig. S5 Proteomic analysis of HEK293 wildtype, XBP1-mN and ATF6-GFP cells treated with Oxaliplatin, Thapsigargin or Staurosporine. (A)** The numbers of significantly down/upregulated proteins (*q* ≤ 0.05) in HEK293 WT, XBP1-mN or ATF6-GFP cells treated with Tg (1 μM, 24 h), compared to untreated condition, and overlaps of those for each cell line. **(B)** The numbers of significantly down/upregulated proteins (*q* ≤ 0.05) in WT cells treated with OxaPt (20 μM, 24 h), Tg (1 μM, 24 h) or Staurosporine (1 μM, 24 h), compared to untreated condition, and overlaps of those for each treatment.

**Fig. S6 Comparison of proteome between different treatment in HEK293 wildtype cells.** Volcano plots of down/upregulated proteins (left panels) and enrichment pathway analysis based on gene ontology terms performed for downregulated proteins (right panels, *q* ≤ 0.05 or ≤ 0.2) in HEK293 WT cells treated with OxaPt (**A**, 20 µM, 24 h), Tg (**B**, 1 µM, 24 h) or Staurosporine (**C**, 1 µM, 24 h), compared to untreated condition. OxaPt, Oxaliplatin; Tg, Thapsigargin.

**Fig. S7 Proteome-based comparison of HEK293 XBP1-mN, ATF6-GFP treated with Thapsigargin**. Volcano plots of down/upregulated proteins (left panels) and enrichment pathway analysis based on gene ontology terms performed for upregulated proteins (right panels, *q* ≤ 0.05 or ≤ 0.2) in **A**) HEK293 XBP1-mN and **B**) ATF6-GFP cells treated with Tg (1 µM, 24 h), compared to untreated condition. Tg, Thapsigargin.
