## Supplementary Figures for "Endoplasmic reticulum stress signaling actively contributes to therapy resistance in colorectal cancer"

Fig.S1

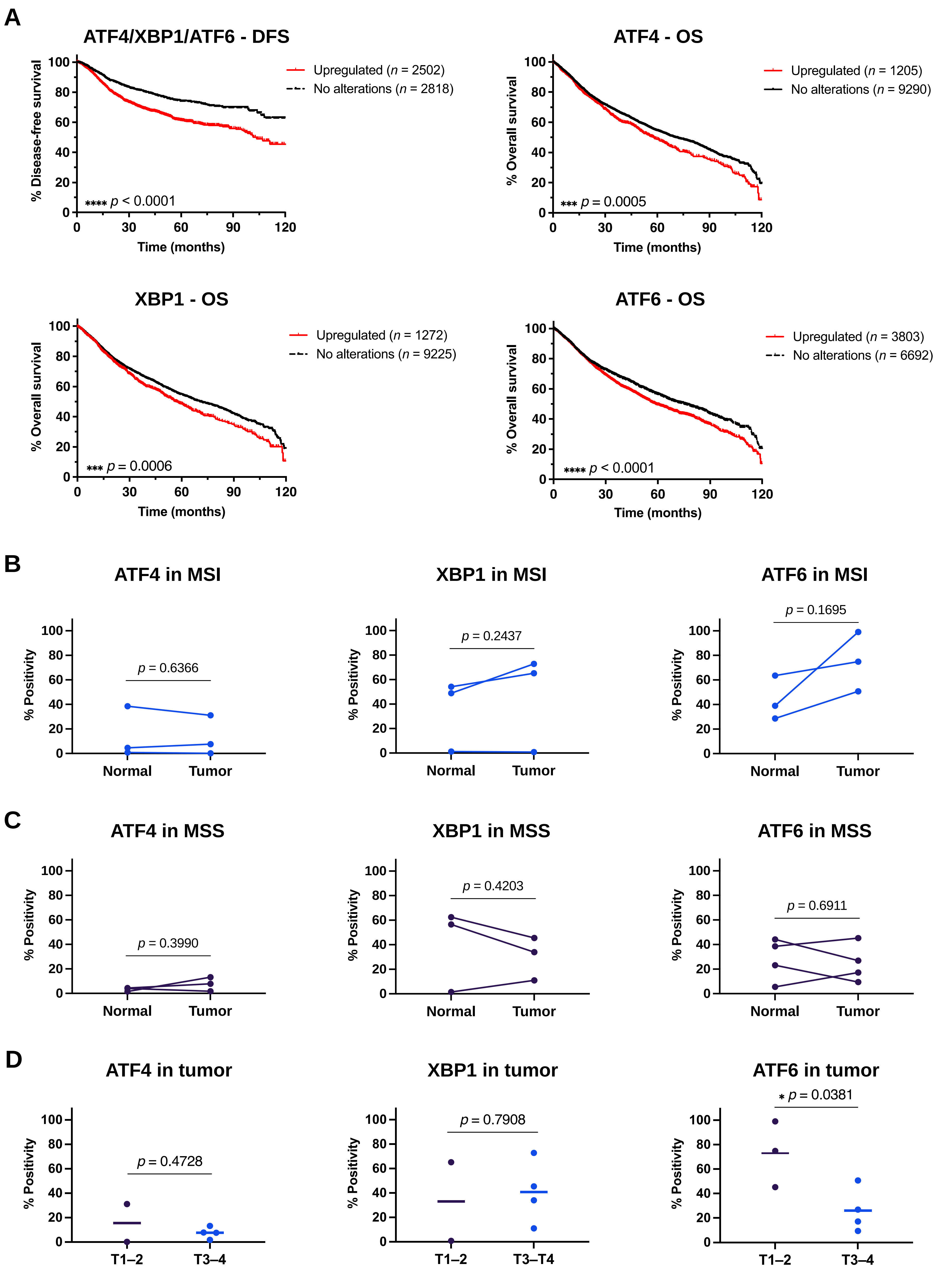
A
ATF4/XBP1/ATF6 - DFS
% Disease-free survival
Upregulated (n = 2502)
No alterations (n = 2818)
**** p < 0.0001
Time (months)
ATF4 - OS
% Overall survival
Upregulated (n = 1205)
No alterations (n = 9290)
*** p = 0.0005
Time (months)
XBP1 - OS
% Overall survival
Upregulated (n = 1272)
No alterations (n = 9225)
*** p = 0.0006
Time (months)
ATF6 - OS
% Overall survival
Upregulated (n = 3803)
No alterations (n = 6692)
**** p < 0.0001
Time (months)
B
ATF4 in MSI
p = 0.6366
XBP1 in MSI
p = 0.2437
ATF6 in MSI
p = 0.1695
% Positivity
Normal
Tumor
C
ATF4 in MSS
p = 0.3990
XBP1 in MSS
p = 0.4203
ATF6 in MSS
p = 0.6911
% Positivity
Normal
Tumor
D
ATF4 in tumor
p = 0.4728
XBP1 in tumor
p = 0.7908
ATF6 in tumor
* p = 0.0381
% Positivity
T1–2
T3–4
T3–T4

Fig.S2

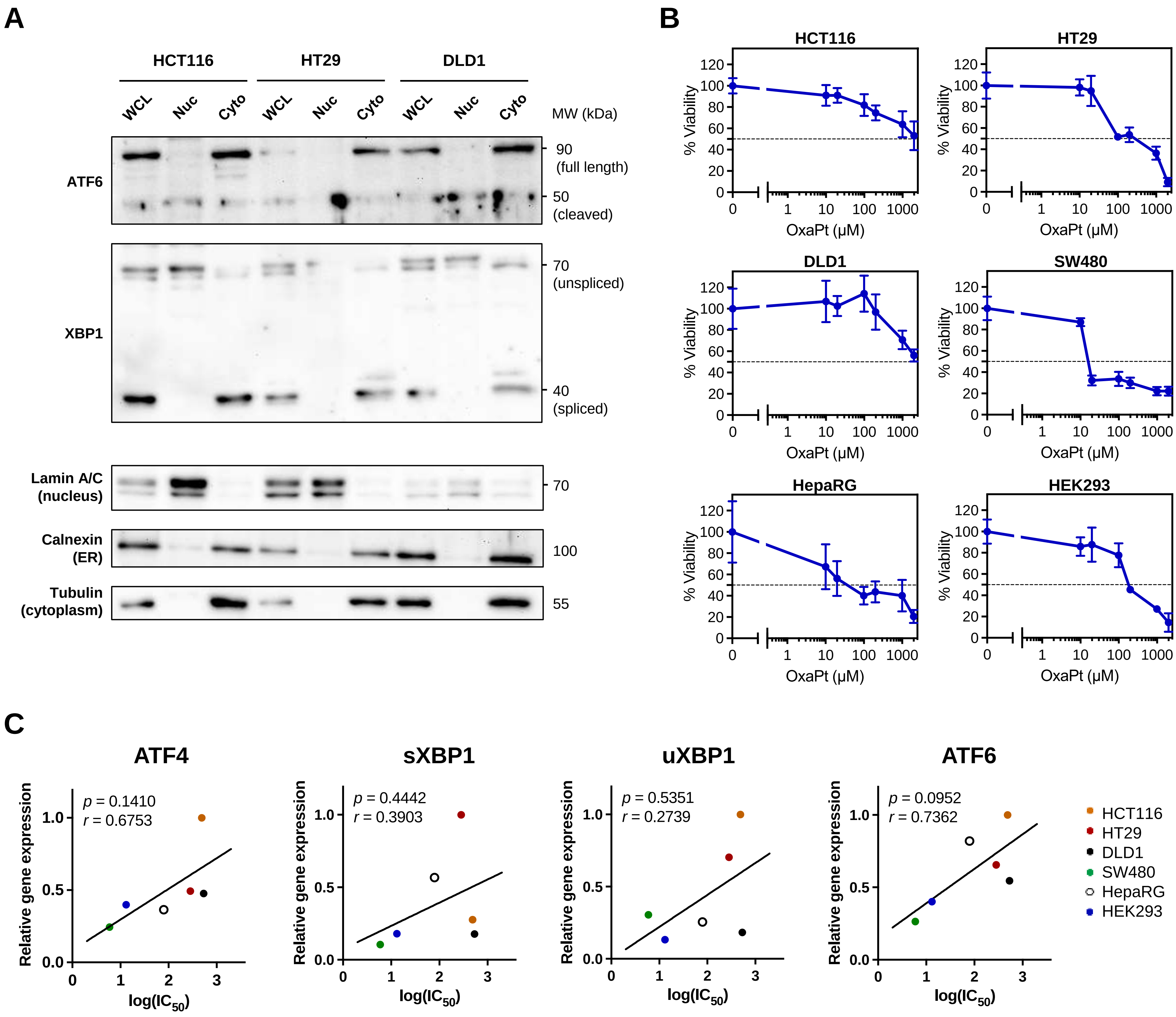
A
HCT116
HT29
DLD1
WCL
Nuc
Cyto
MW (kDa)
ATF6
90
(full length)
50
(cleaved)
XBP1
70
(unspliced)
40
(spliced)
Lamin A/C
(nucleus)
70
Calnexin
(ER)
100
Tubulin
(cytoplasm)
55
B
HCT116
HT29
DLD1
SW480
HepaRG
HEK293
% Viability
OxaPt (μM)
C
ATF4
p = 0.1410
r = 0.6753
sXBP1
p = 0.4442
r = 0.3903
uXBP1
p = 0.5351
r = 0.2739
ATF6
p = 0.0952
r = 0.7362
Relative gene expression
log(IC50)
HCT116
HT29
DLD1
SW480
HepaRG
HEK293

**Fig.S3**

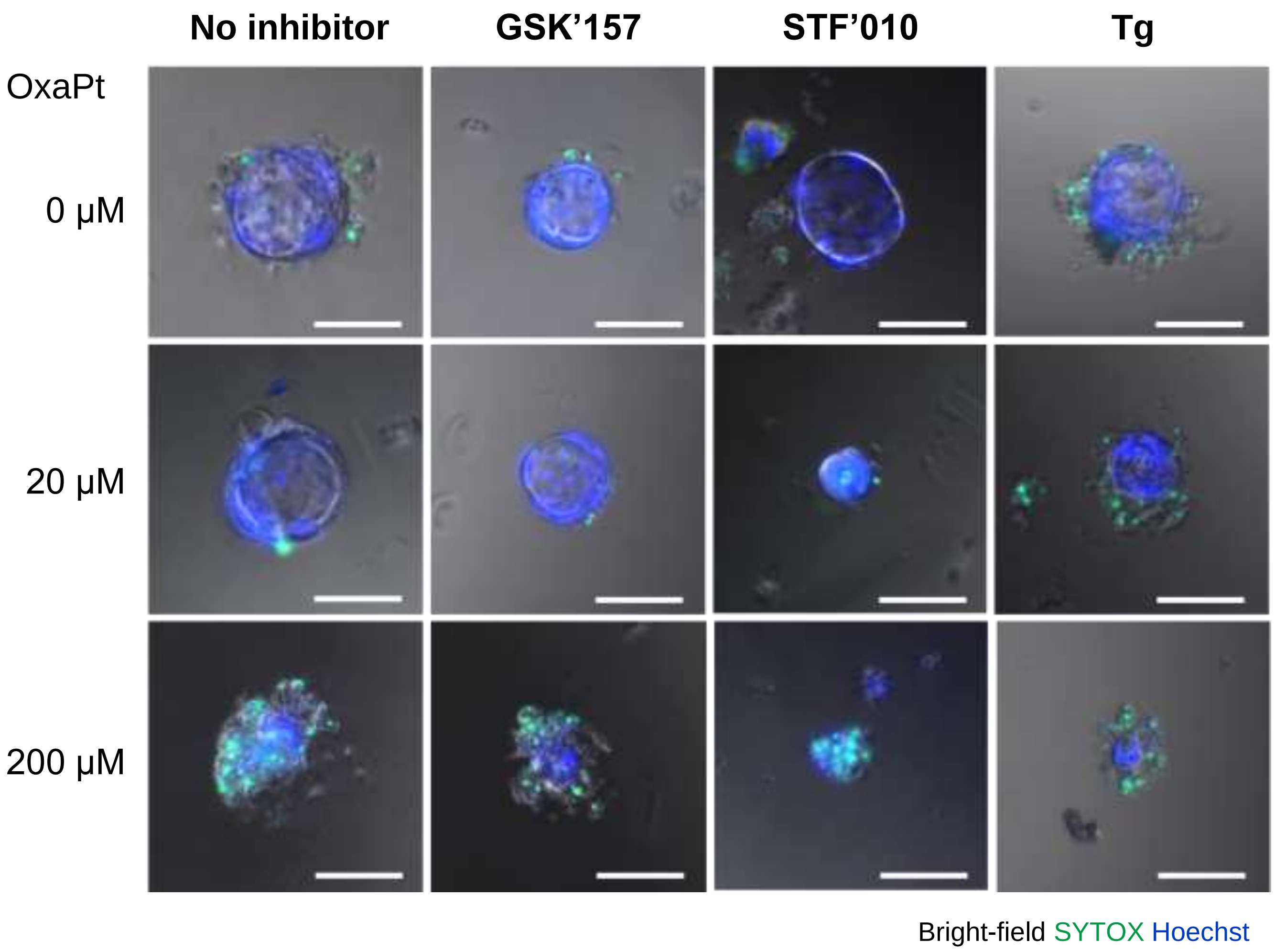
No inhibitor
GSK'157
STF'010
Tg
OxaPt
0 μM
20 μM
200 μM
Bright-field SYTOX Hoechst

Fig.S4

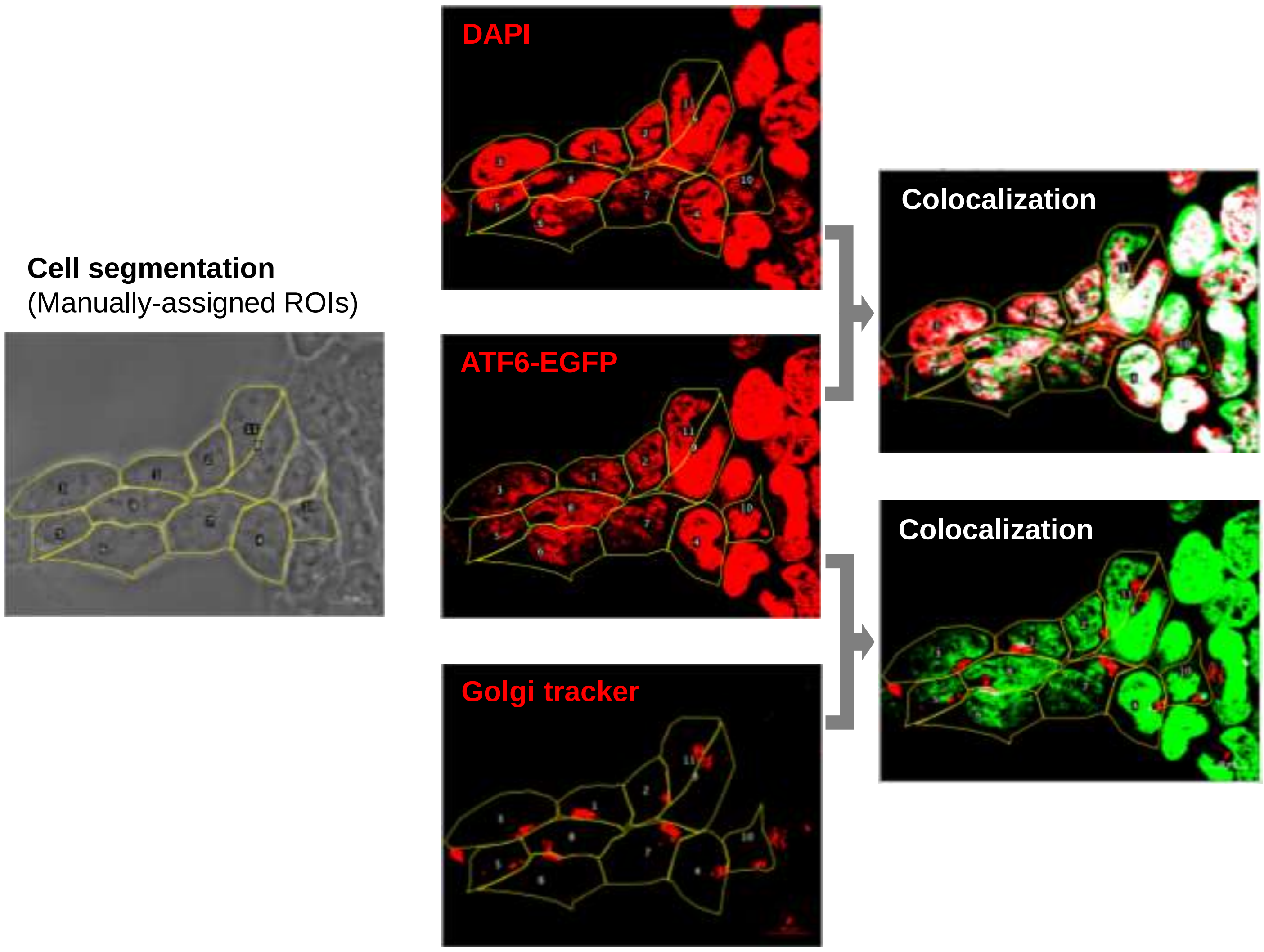


Data analysis with ImageJ software (ROI drawing, thresholding, colocalization, pixel counting).
Plugin: Colocalization (https://github.com/Anders-Lunde/Colocalization_Object_Counter)

#### Fig.S5

**A**

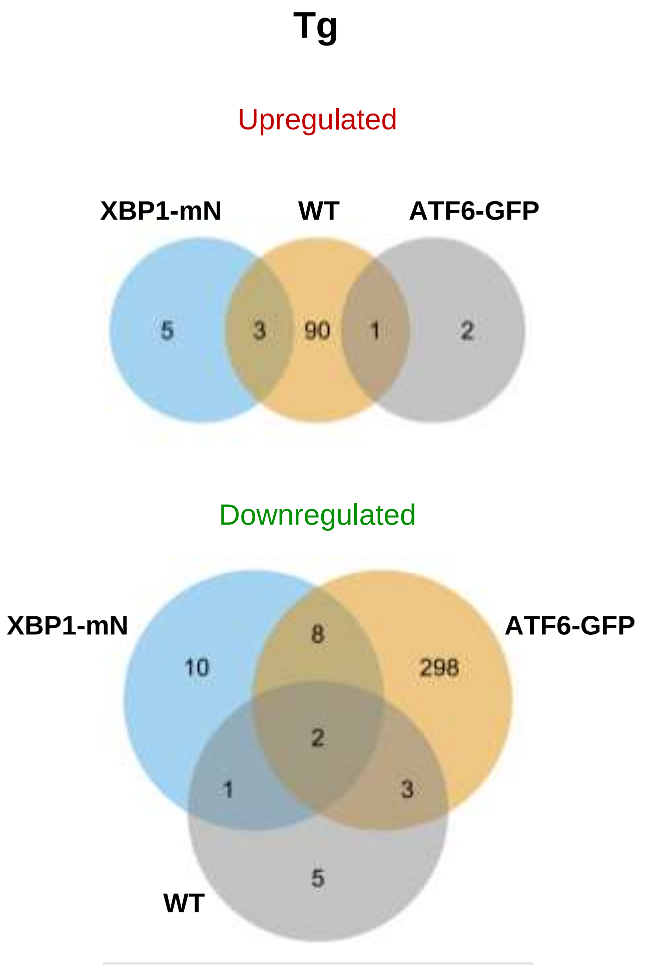
Tg
Upregulated
XBP1-mN
WT
ATF6-GFP
5
3
90
1
2
Downregulated
XBP1-mN
ATF6-GFP
10
8
298
2
1
3
5
WT

**B**

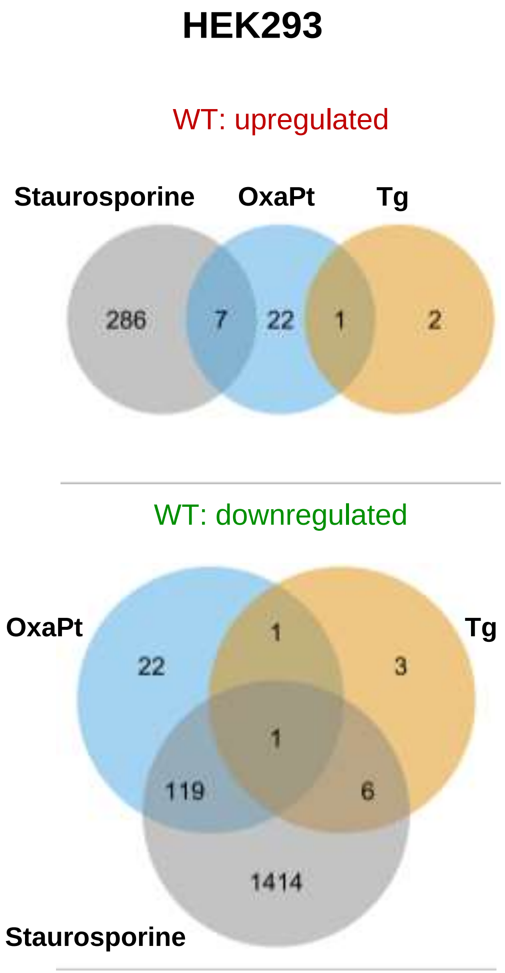
HEK293
WT: upregulated
Staurosporine
OxaPt
Tg
286
7
22
1
2
WT: downregulated
OxaPt
Tg
22
1
3
1
119
6
1414
Staurosporine

### Fig. S6

## A

##### HEK293 WT - OxaPt vs. untreated

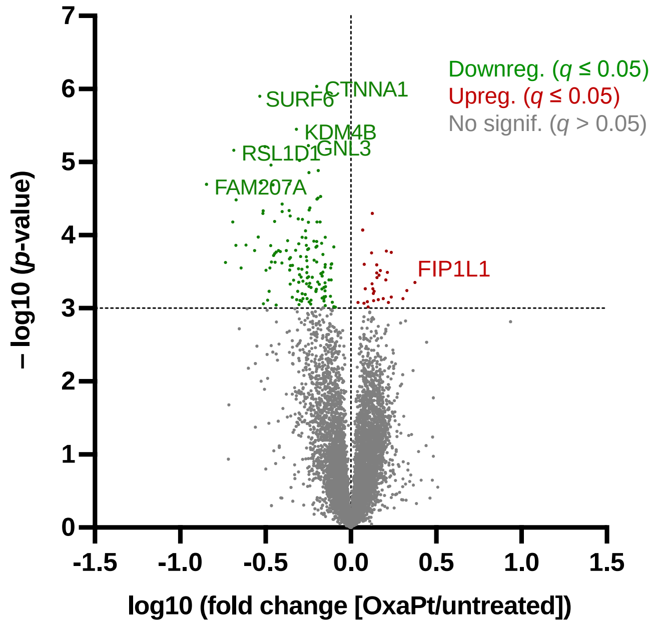


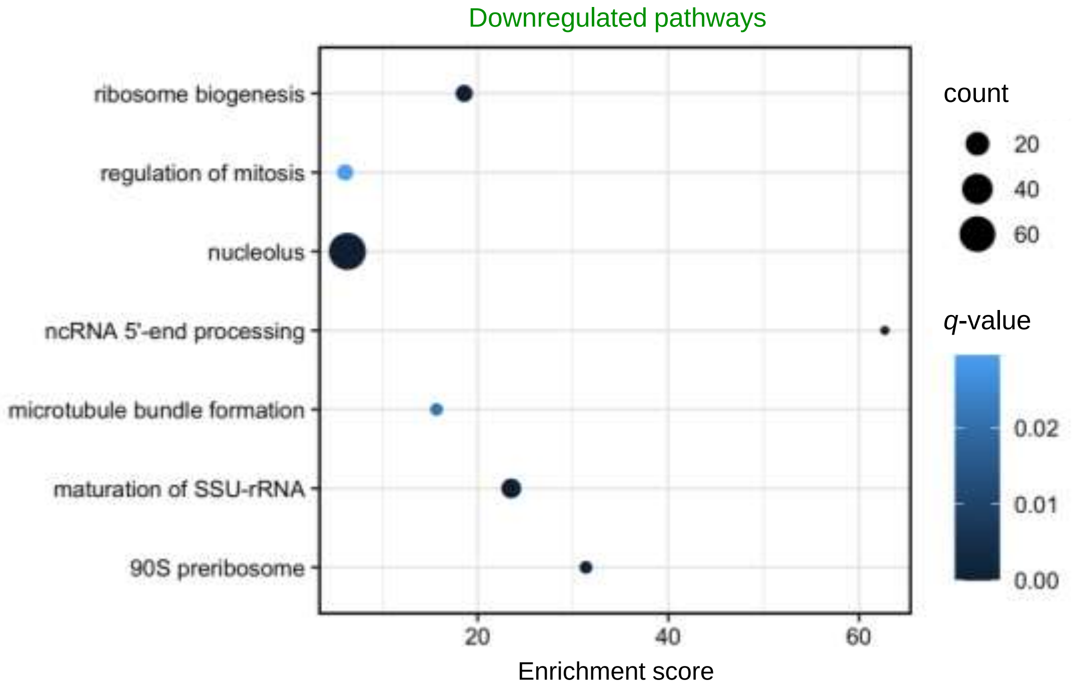


## B

##### HEK293 WT - Tg vs. untreated

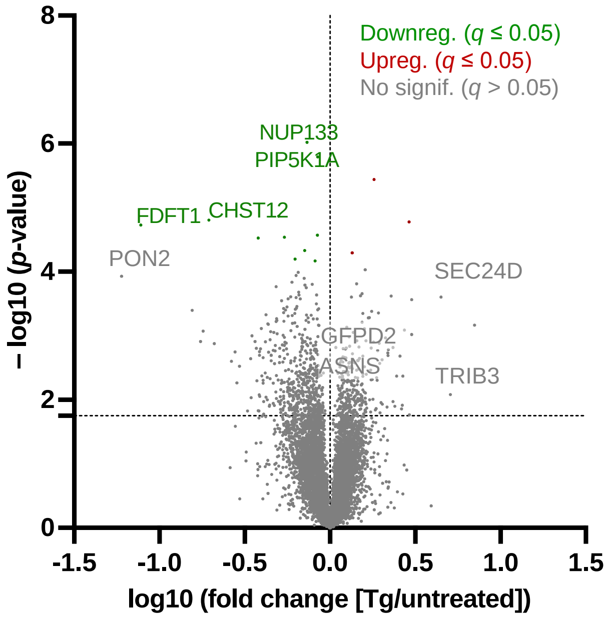


No apparent changes on the global proteome
($q \le 0.2$)

## C

##### HEK293 WT - Staurosporine vs. untreated

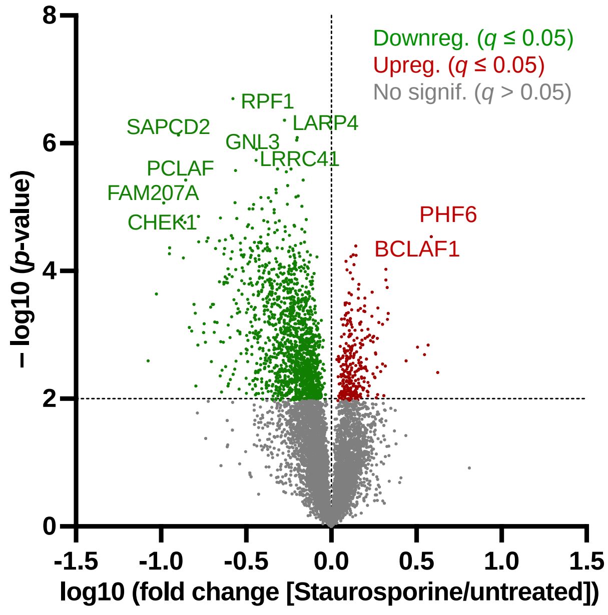


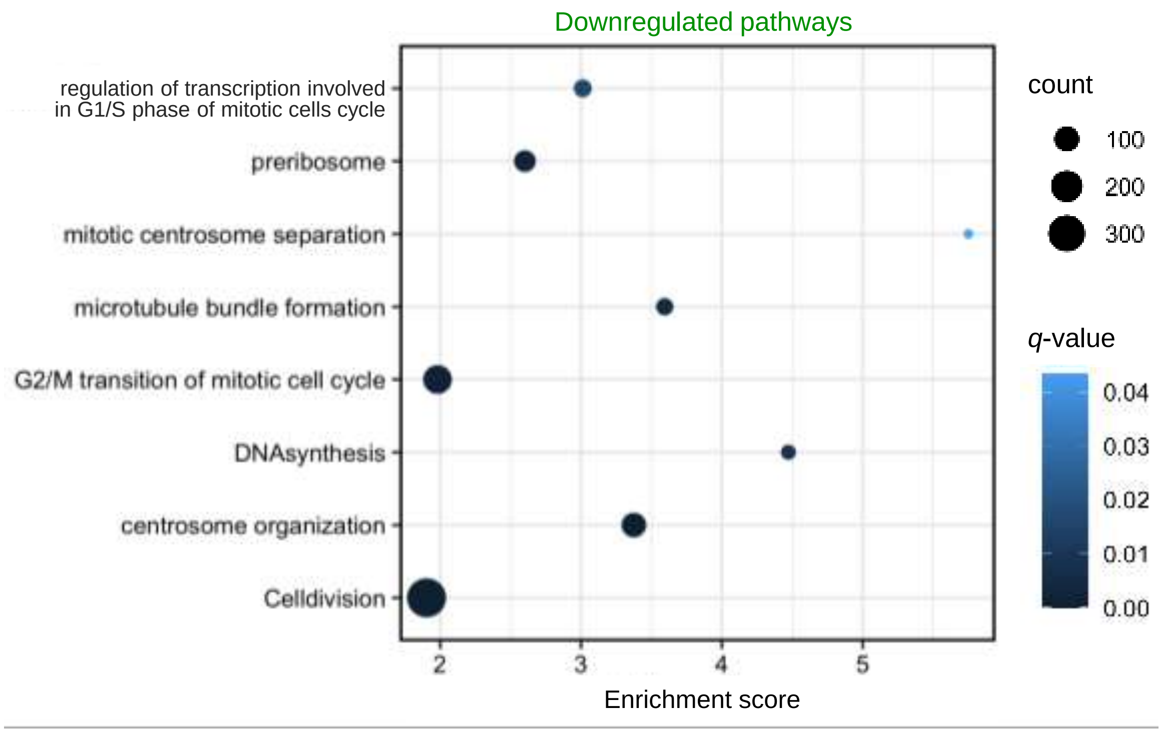

**Fig. S7**

**A**

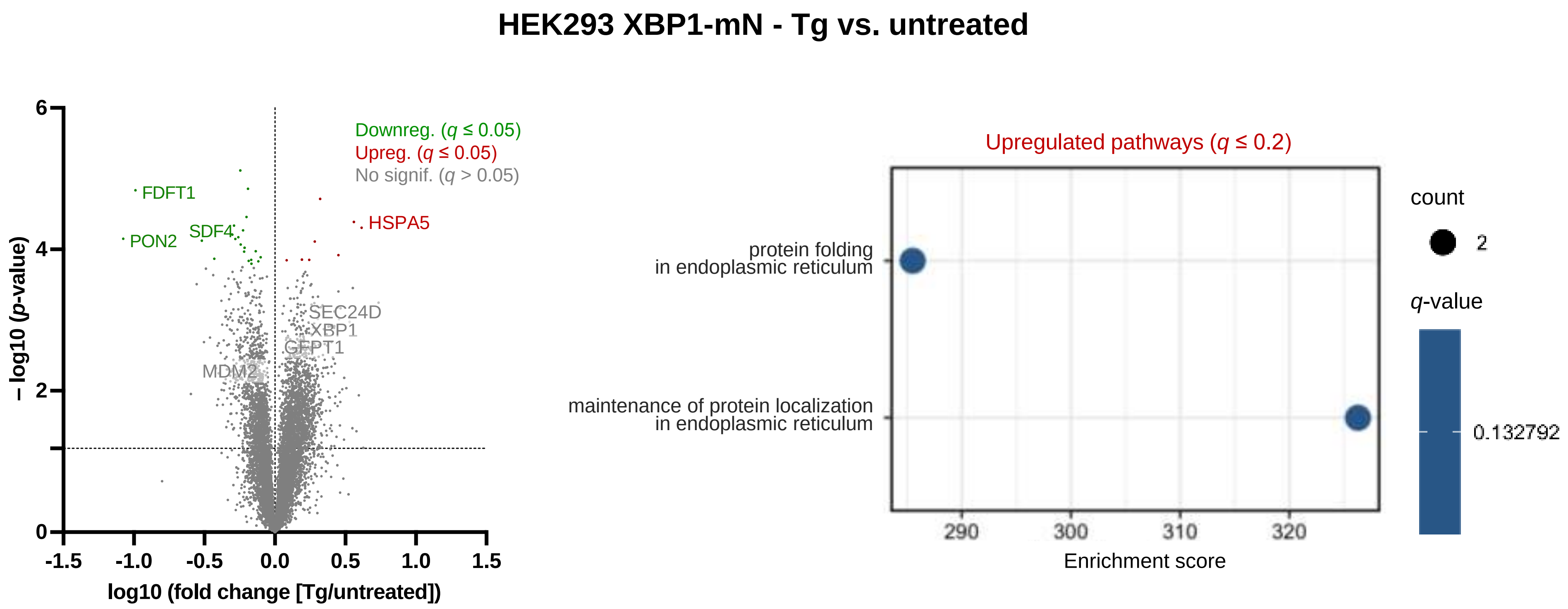
HEK293 XBP1-mN - Tg vs. untreated
Downreg. (q ≤ 0.05)
Upreg. (q ≤ 0.05)
No signif. (q > 0.05)
FDFT1
PON2
SDF4
HSPA5
SEC24D
XBP1
GFPT1
MDM2
− log10 (p-value)
log10 (fold change [Tg/untreated])
Upregulated pathways (q ≤ 0.2)
protein folding
in endoplasmic reticulum
maintenance of protein localization
in endoplasmic reticulum
Enrichment score
count
q-value
0.132792

**B**

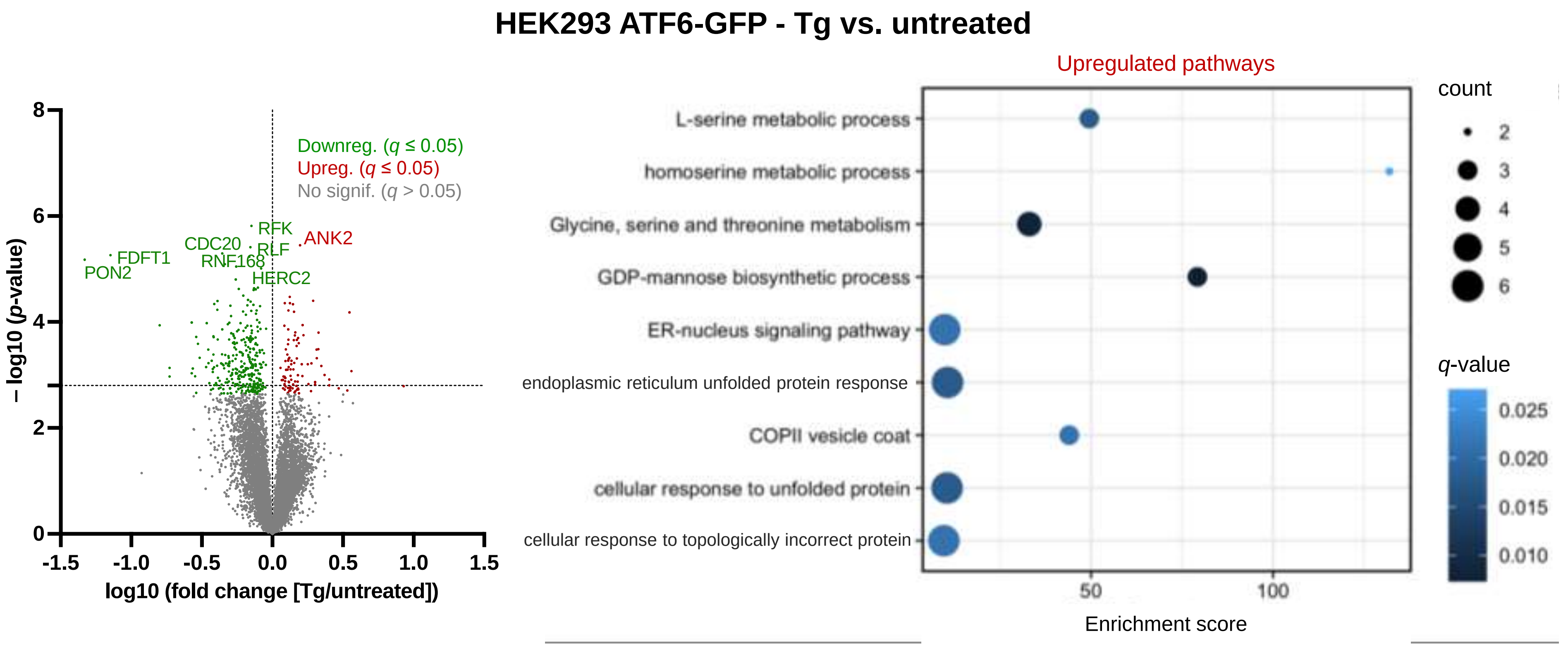
HEK293 ATF6-GFP - Tg vs. untreated
Downreg. (q ≤ 0.05)
Upreg. (q ≤ 0.05)
No signif. (q > 0.05)
RFK
ANK2
CDC20
RLF
FDFT1
RNF168
PON2
HERC2
− log10 (p-value)
log10 (fold change [Tg/untreated])
Upregulated pathways
L-serine metabolic process
homoserine metabolic process
Glycine, serine and threonine metabolism
GDP-mannose biosynthetic process
ER-nucleus signaling pathway
endoplasmic reticulum unfolded protein response
COPII vesicle coat
cellular response to unfolded protein
cellular response to topologically incorrect protein
Enrichment score
count
q-value
